# Contractile ring composition dictates kinetics of *in silico* contractility

**DOI:** 10.1101/2022.07.06.499011

**Authors:** Daniel B. Cortes, Francois Nedelec, Amy Shaub Maddox

## Abstract

Constriction kinetics of the cytokinetic ring are expected to depend on dynamic adjustment of ring composition, but the impact of component abundance dynamics on ring constriction is understudied. Computational models generally assume that contractile networks maintain constant composition. To test how compositional dynamics affect constriction kinetics, we first measured F-actin, non-muscle myosin II, septin, and anillin during *C. elegans* zygotic mitosis. A custom microfluidic device that positioned the cell with the division plane parallel to a light sheet allowed even illumination of the cytokinetic ring. Measured component abundances were implemented in an agent-based 3D model of a membrane-associated contractile ring. With constant network composition, constriction occurred with biologically unrealistic kinetics. Measured changes in component quantities elicited realistic constriction kinetics. Simulated networks were more sensitive to changes in motor and filament amounts, than that of crosslinkers and tethers. Our findings highlight the importance of network composition for actomyosin contraction kinetics.

**Summary:** We created a microfluidic device compatible with high numerical aperture light sheet microscopy to measure cytokinetic ring component abundance in the C. elegans zygote. Implementing measured dynamics into our three-dimensional agent-based model of a contractile ring elicited biologically realistic kinetics.

## Introduction

In animal cell cytokinesis, one cell is physically separated into two by a plasma membrane furrow drawn in by a dynamic cytoskeletal contractile ring. The contractile ring is rich in actin filaments (F-actin), non-muscle myosin II (NMMII) filaments, and other structural components including non-motoring crosslinkers like anillin, and other scaffold proteins including septins. The anaphase spindle initiates a Rho signaling cascade that concentrates contractile ring components at the equatorial cortex (Bement et al., 2005; Nishimura and Yonemura, 2006; Pollard and O’Shaughnessy, 2019) first as a loose isotropic mesh that reorganizes into a tight circumferentially aligned cord, which closes the cytoplasmic connection between daughter cells (White and Borisy, 1983; Reymann et al., 2016; Descovich et al., 2018; Sobral et al., 2021). The two nascent cells become topologically distinct when the intercellular midbody bridge is severed during abscission. Accurate genomic inheritance requires complex regulation of cytokinesis, from initial placement of the assembling actomyosin structure (Werner et al., 2007; Basant and Glotzer, 2018), to proper regulation of constriction dynamics and irreversible completion (Bembenek et al., 2013; Petsalaki and Zachos, 2021). As the contractile ring constricts, the abundance of structural components changes due to continued recruitment, compaction, disassembly, depolymerization, inhibition, and other processes. Early electron microscopic studies of cytokinetic rings suggested that the overall density of the cortical cytoskeleton was constant throughout ring closure (Schroeder, 1972). More recently, changes in the density of several contractile ring components have been reported: for NMMII (Carvalho et al., 2009; Wu and Pollard, 2005; Khaliullin et al., 2018; Wollrab et al., 2016; Xue and Sokac, 2016), F-actin (Wu and Pollard, 2005; Xue and Sokac, 2016; Wollrab et al., 2016), septin (Carvalho et al., 2009; Xue and Sokac, 2016), and anillin (Wu and Pollard, 2005; Carvalho et al., 2009; Khaliullin et al., 2018). In the fission yeast *S. pombe*, the density of these, and other, components first increases as the contractile ring matures and then decreases in the latter half of constriction (Wu and Pollard, 2005). In *C. elegans* blastomere cytokinesis, though changes are less pronounced, an increase in overall density is visible late in division (Carvalho et al., 2009), and in zygotic division evidence exists for exponential increase in density of both NMMII and anillin throughout cytokinesis (Khaliullin et al., 2018). These studies suggest that the overall composition of the contractile ring can change throughout constriction, with components exhibiting differential changes in density that would result in different ratios between them (Wu and Pollard, 2005; Carvalho et al., 2009; Xue and Sokac, 2016). However, in some cases, multiple contractile ring components exhibit parallel changes in abundance, suggesting consistent composition and the potential for contractile units (Khaliullin et al., 2018). Dynamic differences in ring composition could account for variations in constriction kinetics and overall cytokinesis timing among different cell types and species. We predicted that, beyond the “parts list” of conserved, essential structural components, the dynamic abundance of these components impacts cytokinetic ring dynamics. Indeed, the abundance of NMMII motors and non-motor cross linkers has a non-linear effect on contractile ring dynamics (Ennomani et al., 2016; Ding et al., 2017; Descovich et al., 2018), further highlighting the importance of contractile ring composition and its dynamics for constriction kinetics.

Computer models are powerful tools for evaluating the potential consequences of measured changes in actomyosin ring composition. Though it comes with its own set of limitations, the ability to model with high spatial and temporal resolution serves as a complimentary approach for cell biology where microscopy can suffer from resolution limits. Two-dimensional agent-based models have successfully captured how changes in crosslinker density affect the contractility of an isotropic network (Belmonte et al., 2017) and isotropic rings (Ennomani et al., 2016; Descovich et al., 2018), but differences between ring organization in vivo and in silico (isotropic versus sarcomeric) (Capco and Bement, 1991; Carvalho et al., 2009) may limit the relevance of these simulations. Agent-based models of cytokinesis have been used extensively to predict and describe physical concepts that drive contractile ring formation (Vavylonis et al., 2008; Sobral et al., 2021) and general mechanisms of actomyosin contraction (Mendes Pinto et al., 2012; Oelz et al., 2015; Thiyagarajan et al., 2017; Nguyen et al., 2018). While these models have yielded insights into universal properties of actomyosin networks including contractile rings, none has considered changes in network composition throughout the process of contraction. We chose the mitotic cytokinetic rings of the *C. elegans* zygote as the biological model for our simulation, because *i)* its evolution is stereotyped and well-characterized, *ii)* its contraction is a cell-autonomous event, *iii)* fluorescent CRISPR transgenes with endogenous expression are readily availability, *iv)* it can be imaged in toto using live single-plane illumination microscopy, v) it can be resistant to phototoxic damage under certain long-term imaging conditions (Rog and Dernburg, 2015). Previous *C. elegans* zygote imaging methods relied upon imaging of the contractile ring in profile view and were subject to reconstruction artifacts, anisotropic resolution, and depth penetration issues. Fission yeast and HeLa cells were previously manipulated into miniature wells (Wollrab et al., 2016) to place the contractile ring parallel to the focal plane, such that confocal acquisition could be reduced to a few z-sections rather than a full cell volume. Such *en face* imaging reduces artifacts and permits higher temporal resolution.

Here, we performed *en face* imaging with a custom built microfluidic chip based on a staging device (Cornaglia et al., 2015), combined with high-resolution light sheet microscopy (Fadero et al., 2018). Further illuminating the contractile ring from the side with a light-sheet, which allowed isotropic spatial resolution and minimal reconstruction or depth artifacts. We measured the dynamic composition of contractile rings, specifically changes of NMMII, F-actin, anillin and septin throughout cytokinesis. We implemented these measurements in a three-dimensional agent-based model that consists of actin-like filaments, NMMII-like motor filaments, non-motor crosslinkers, and tethers that link the network to a deformable surface that represents the plasma membrane. This model revealed that constriction kinetics better reflect measured biological kinetics when composition changes are considered. With constant composition, simulations failed to recapitulate our biological measurements of cytokinetic kinetics. We further identify specific contractile ring components, NMMII and F-actin filaments as particularly important for dictating constriction kinetics.

## Results

### Abundance dynamics measurements from C. elegans zygotic division

The compositional dynamics of contractile rings have been measured for some components in some cell types (Wu and Pollard, 2005; Carvalho et al., 2009; Wollrab et al., 2016; Khaliullin et al., 2018), but a systematic study has not been undertaken in an animal cell. We therefore first measured contractile ring composition changes throughout *C. elegans* zygotic division. To generate data with high spatiotemporal resolution and with minimal reconstruction or imaging artifacts, we imaged the zygotic contractile ring *en face* using custom-made PDMS microfluidic devices, which were designed based on similar devices that immobilize *C. elegans* embryos in a horizontal orientation (Cornaglia et al., 2015). Our device trapped dissected zygotes in an upright position and allowed us to illuminate the contractile ring with a Mizar TILT lateral-excitation light sheet system (Fadero et al., 2018). We imaged cytokinesis in zygotes expressing endogenously-expressed fluorescent markers for non-muscle myosin II (NMMII; NMY-2::GFP; Movie 1), F-actin (mKate::LifeAct; Movie 2), anillin (mNeonGreen::ANI-1; Movie 3), and septin (UNC-59::GFP; Movie 4). Z-stacks were acquired from the onset of furrowing through the end of constriction. Maximum intensity projections through the thickness of the contractile ring revealed differences among markers for normalized ring intensity dynamics throughout cytokinesis (Figure 1B-E, B’-E’). NMMII (n = 10) and F-actin (n = 12) markers increased in normalized intensity in a linear fashion as cytokinesis progressed (Figure 1B’, C’). Septin (n = 15) and anillin (n = 14), on the other hand, increased exponentially in normalized intensity (Figure 1D’, E’). Normalized intensity measurements provided an estimate of the relative density changes for each of the four measured contractile ring components (Figure 1B-E”). Relative density changes described composition changes per unit area of the contractile ring as opposed to describing changes in whole contractile ring composition. Thus, we multiplied the density curves by contractile ring perimeter (which was derived from radius measurements) to estimate total fold change of each contractile ring component (Figure S3A-D, A’-D’). Since the actual amount of any component in the *C. elegans* zygote cytokinetic ring has not been measured, we estimated maximum protein amounts for each component using measurements of total amounts on the *S. pombe* contractile ring (Wu and Pollard, 2005; Courtemanche et al., 2017). For each ring component, we normalized our estimated fold-change to these maximum values, resulting in an estimated amount of each ring component throughout cytokinesis (Figure S3A-D’) in a *C. elegans* embryo-like cell the size of a *S. pombe* cell. These data were then incorporated into our Cytosim simulations of deformable ring space as a function of space radius, to match the total amount of each of the four simulated agents representing the measured contractile ring components to the measurements for the corresponding component *in vivo* at every simulated time point (Figure S4F-G).

**Figure 1.**
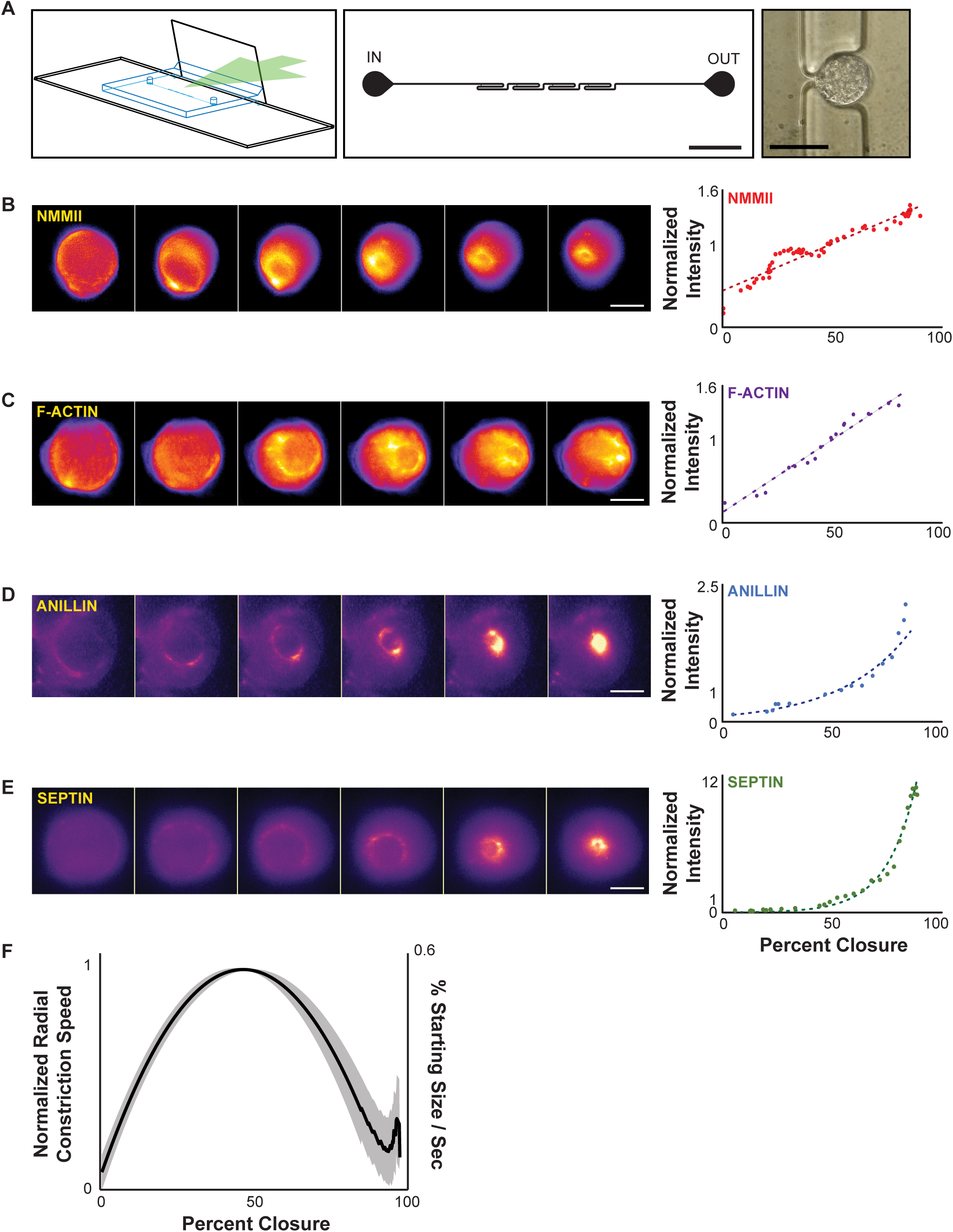
Live imaging *of C. elegans* zygotic contractile rings. A) Left: Graphical representation of custom microfluidic trap on a 25 x 60 mm coverslip (black) with a second coverslip approximately perpendicular to the imaging plane (black; green arrow). Center: Microfluidic trap design showing the inlet and outlet channels and four trap features; scale bar = 2 mm. Right: Representative *C. elegans* embryo standing upright in trap cup; scale bar = 30 μm. B) Left: Time lapse series of the contractile ring in a *C. elegans* zygote expressing fluorescently tagged NMMII (NMY-2::GFP; Movie 1). Right: Normalized mean pixel intensity plot of the contractile ring of a representative cell expressing NMY-2::GFP. Dotted red line: best fit line. n = 10. C) Left: Time lapse series of contractile ring in a *C. elegans* zygote expressing a fluorescent F-actin reporter transgene (mKate::LifeAct; Movie 2). Right: Normalized mean pixel intensity plot of the contractile ring of a representative cell expressing mKate::LifeAct. Dotted purple line: best fit line. n = 12. D) Left: Time lapse series of contractile ring in a *C. elegans* zygote expressing a fluorescently-tagged anillin (mNeonGreen::ANI-1; Movie 3). Right: Normalized mean pixel intensity plot of the contractile ring of a representative cell expressing mNeonGreen::ANI-1. Dotted blue line: best fit exponential curve. n = 14. E) Left: Time lapse series of contractile ring in a *C. elegans* zygote expressing a fluorescently-tagged septin (UNC-59::GFP; Movie 4). Right: Normalized mean pixel intensity plot of the contractile ring of a representative cell expressing UNC-59::GFP. Dotted green line: best fit exponential curve. n = 15. F) Plot of cytokinetic ring constriction speed (originally peaking at 0.54% starting ring size per second; normalized so maximum speed is 1.0) in the *C. elegans* zygote. Gray area around the solid black line: mean ± standard deviation. n = 23. Scale bars = 15 μm.

### Multi-dimensional analysis of C. elegans contraction kinetics

In order to compare *in vitro* and *in silico* closure kinetics, we next pooled the imaging data together to estimate the average closure kinetics curve for our biological dataset (Figure 1F, B, n = 27). We characterized ring closure with five metrics: normalized maximum speed, percent closure at maximum speed, peak acceleration, peak deceleration, and the ratio between acceleration and deceleration (as a measure of the symmetry of the kinetics curve). These measurements of our biological data were assembled into a normalized pentagon wherein the average value of each parameter *in vivo* was set as the standard (i.e 1.0 for each; Figure S1E). Zygote ring contraction speed increased until 47% closure (Figure S1E; Table S1), at which point it reached approximately 0.5%/sec (normalized to 1.0 below). Acceleration and deceleration rates were indistinguishable at 7.9×10^-3^ %/sec^2^); the ratio of acceleration to deceleration was 1.0 (Table S1). The area of the pentagon served as a composite score to semi-quantitatively compare conditions, and the shape of the pentagon offers a qualitative representation of the five metrics (Figure S1E).

### Simulated contractile rings with C. elegans abundance dynamics recapitulate biological constriction kinetics

Generic simulations of contractile rings incorporate actin-like filaments, various configurations of motors, and non-motor crosslinkers (Mendes Pinto et al., 2012; Oelz et al., 2015; Ennomani et al., 2016). However, these components are generally present at some user specified amount that does not vary throughout the simulated constriction, completely ignoring composition changes. To test the impact of measured changes in ring composition on contraction kinetics, we simulated a 3-dimensional contractile network tethered to a deformable space (Figure 2A, B, Figure S2A). Such models have been implemented successfully for fission yeast cytokinesis (Nguyen et al., 2018), but without implementation of protein abundance dynamics. In this work adapted a 3-dimensional deformable space in Cytosim to act as a semi-rigid proxy to the plasma membrane (Dmitrieff et al., 2017) and used a simple geometric approximation of cell deformation throughout cytokinesis to estimate a surface tension-based resistance to deformation of the membrane (Sain et al., 2015). Within this framework we further added an algorithm to change contractile ring component abundance in real time to match total protein amounts estimated from measured density changes in live cells (Figure S3, Figure S4). Our algorithm removed components from the simulation space or added them proximal to the deformable surface at each time point, according to instantaneous contractile ring size (see MM2). In this way, we simulated the changes in the composition of contractile ring components (Figure S4A’-D’). Furthermore, we implemented a complex method for simulating each of the four contractile ring components as filaments, made up of points connected by springs, with permanently attached “domains” that can bind other components. In this way, we simulated NMMII-like motor ensemble with 30 discrete filament binding motors in a bipolar arrangement. This approach also allowed us to simulate anillin-like crosslinkers with multiple types of binding interactions as anillin has been shown to interact with both F-actin and NMMII filaments (Straight et al., 2005).

**Figure 2.**
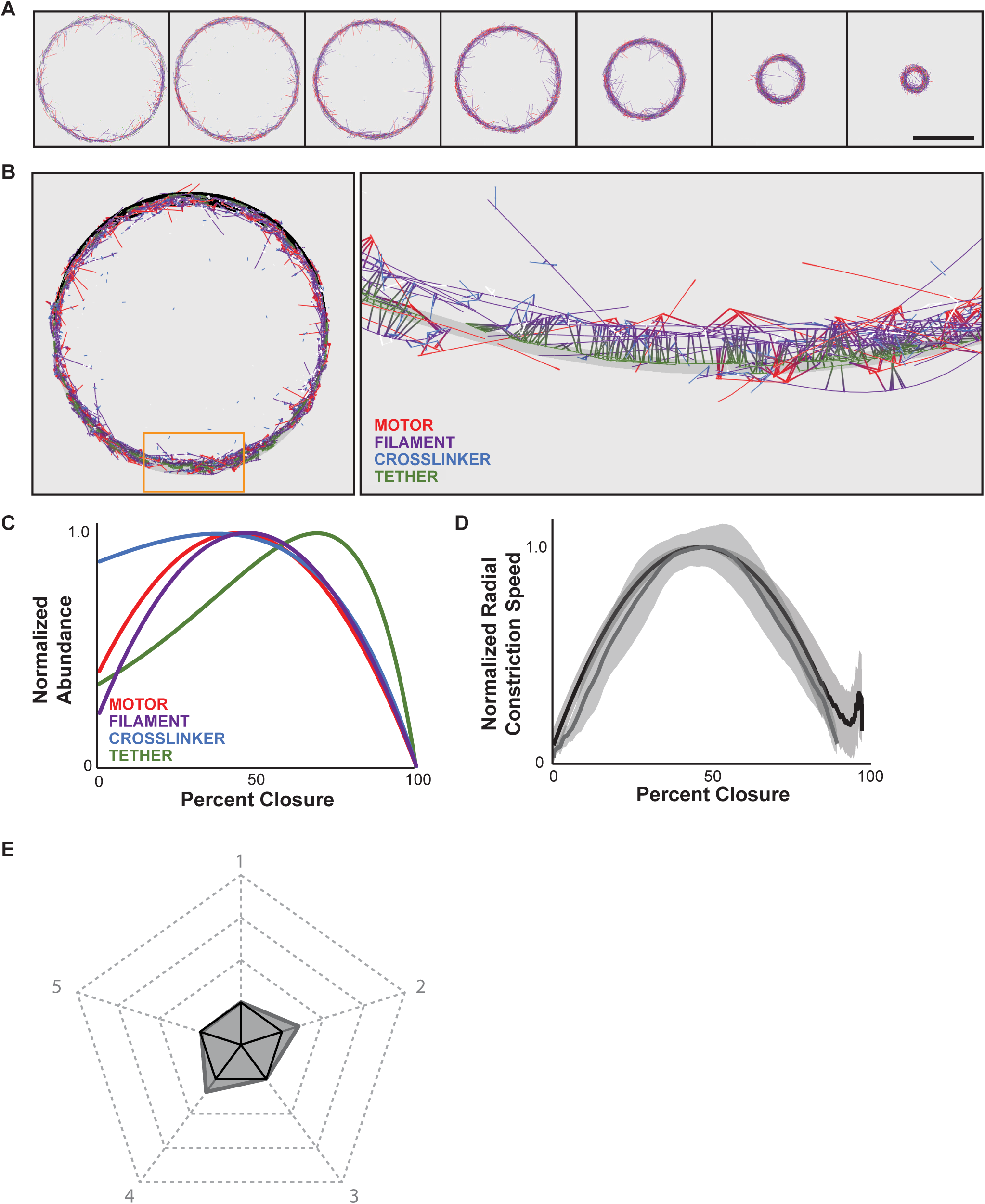
Simulated contractile rings with biologically-informed abundance dynamics exhibit realistic closure kinetics. A) Time series from a representative three-dimensional simulation viewed as a two-dimensional projection, showing the four main simulated contractile ring components. Dynamic space is tethered to the contractile ring trailing (outside) edge. Same simulation as shown in Movie 5 and Movie 6. B) Left: Full view of a representative simulated contractile ring with dynamic ring space shown in black. Right: Magnified view of orange-rectangle region in A; dynamic space is shown as a gray surface. Tether components (green) are tightly associated with the gray surface. All other components associate with the deformable space indirectly through association with the tethering component, except for the filaments (purple) whose plus ends are also tethered directly to the surface. C) Normalized abundance of four main simulated contractile ring components as a function of constriction of the dynamic space. D) Average constriction dynamics curve of simulated contractile ring (gray line) and *C. elegans* zygotic cytokinetic ring (black line). Gray areas: mean ± standard deviation. n = 30. E) Pentagon generated from the five contractile metrics measured (gray) compared to the biological standard metrics (black solid pentagon). Scale bar = 15 μm.

We first tested the effects of modulating all four major contractile ring components (Figure 2A, C; Movie 5). All simulations with biological abundance dynamics closed completely with reproducible kinetics (30/30; Figure 2A, D; Movie 6). Ring closure kinetics quantitatively resembled our cell biological measurements. To compare simulated and biological kinetics, we normalized the average closure curve so that the normalized maximum contractile speed was 1.00. This maximum speed occurred at 46% closure, which was statistically indistinguishable from the biological standard (Table S1). Peak acceleration and deceleration were comparable, at 1.1×10^-2^ %/sec^2^, and the ratio between them was 1.0 (Figure 2D, E, Table S1). Comparing these five kinetics metrics via our pentagon plot revealed a 30% higher composite score for these simulations, compared to our biological data (Figure 2E, Table S1). The increase in composite score was attributed mostly to higher peak acceleration and deceleration of the simulated contractile rings. However, the ratio between these two, along with the normalized maximum speed, and the point at which maximum speed occur were all strikingly similar to biological measurements.

It was not clear whether the prescribed component dynamics were responsible for biologically realistic ring closure kinetics or whether it was simply important that the correct abundance of components was reached at some point in the simulations. To test the latter hypothesis, we next simulated ring closure with component abundance invariably set to the maximum amount calculated from biological data (Figure 3B). In this condition, simulated rings consistently closed beyond 90% (30/30, Figure 3A, C; Movie 7). However, these rings lacked resemblance to biological constriction kinetics. First, for simulations with constant component abundance, normalized maximum speed was significantly higher than for simulations with all component dynamics (the “simulation standard”) and the biological standard, at 1.20 (which would translate to approximately 0.6%/sec). Normalized maximum speed was reached significantly earlier than either standard as well, at 27% closure. Acceleration and deceleration were also higher than in the biological dataset, at 3.3×10^-2^ %/sec^2^ and 1.1×10^-2^ %/sec^2^, respectively; the ratio between these two metrics was also higher than with the biological data at 3.0 (Figure 3C, D; Table S1). Overall, these changes resulted in a 320% increase in the area of the parameter pentagon compared to the biological standard (Figure 3D, Table S1). Altogether, this suggested that while simulations with constant component abundance could generate robust constriction, abundance dynamics were important for recapitulating the constriction kinetics of *C. elegans* zygote cytokinesis.

**Figure 3.**
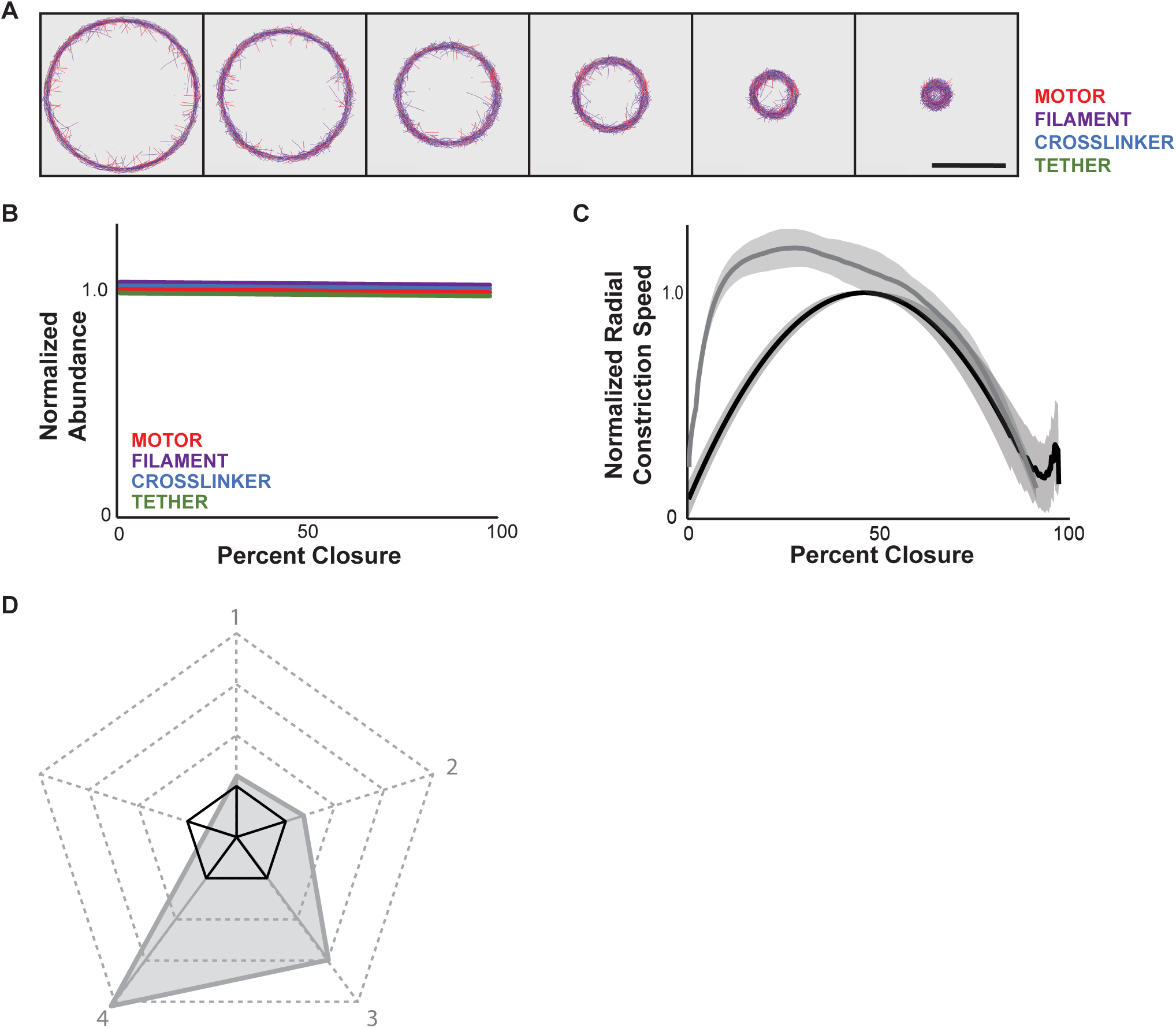
Simulated contractile rings with no abundance dynamics do not exhibit realistic kinetics. A) Time series from a representative three-dimensional simulation viewed as a two-dimensional projection, showing the four main components of the simulated contractile ring and dynamic space. Same simulation as shown in Movie 7. B) Normalized abundance over closure for all four components (constant throughout closure). C) Plot of average constriction dynamics curve of simulated division (gray line) versus the curve from the biological dataset (black line). Gray areas: mean ± standard deviation. n = 30. D) Pentagon generated from the five contractile metrics measured (gray) compared to the biological standard metrics (black solid pentagon). Scale bar = 15 μm.

### The dynamics of motor and filament composition are important for biologically realistic constriction kinetics

While all four contractile ring components simulated here are phylogenetically conserved and essential for normal contractile kinetics in vivo, the effects of their loss of function vary quantitatively. NMMII motoring is inconsistently necessary for successful constriction (Ma et al., 2012; Osorio et al., 2019), and depletion of actin regulator Arp2/3, anillin or septin all result in successful but uncharacteristically concentric division of the *C. elegans* zygote (Dorn et al., 2016). Thus, we next explored how changing composition of each contractile ring component individually affected constriction kinetics. We simulated rings in which the abundances of all components, but one, were set to the maximal amounts, while the altered component was allowed to change in abundance to match biological data (Figure 4A-D).

**Figure 4.**
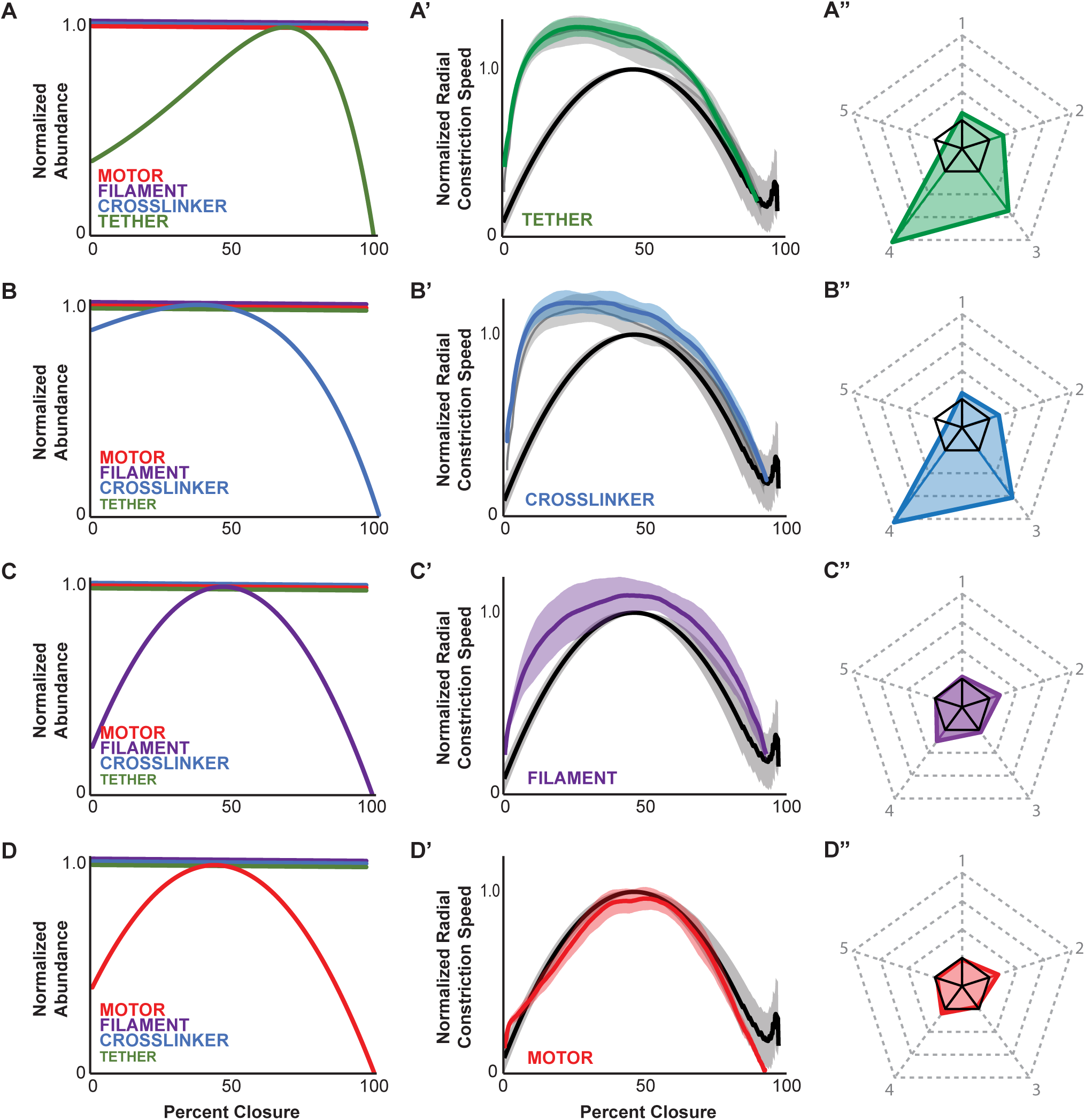
Implementing abundance dynamics of different ring components has distinct effects on constriction. A-D) Normalized component abundance over closure input for simulations with conditions in which each of the four main simulated ring components was individually dynamic. A) Tether only dynamic. B) Non-motor crosslinker only dynamic. C) F-actin-like filament only dynamic. D) Motor ensemble only dynamic. A’-D’) Plots of average constriction dynamics curves of simulated divisions (correspondingly colored curves) versus the curve from the biological dataset (black line; A’-D’) and the simulated data set with no change in composition (light gray in A’, B’). Lighter colored areas around solid curves: mean ± standard deviation. A”-D”) Pentagons generated from the five contractile metrics measured from each simulation setup (color-matched) compared to the biological standard metrics (black solid pentagon). n = 30 for all.

We began with adjustment of the tether component, representing septin in our model, alone (Figure 4A). These contractile rings constricted the dynamic space fully (30/30) but did not exhibit biological kinetics (Figure 4A’). The average constriction curve for these simulations was very similar to that of simulations with no composition changes (Figure 4A’); the parameter pentagon was similarly 310% larger than the biological standard (Figure 4A”, Table S1). Specifically, normalized maximum speed was significantly higher than in the biological and simulation standards, at 1.26, and occurred abnormally early, at 28% closure. Acceleration and deceleration were likewise higher at 3.3×10^-2^ %/sec^2^ and 1.2×10^-2^ %/sec^2^ respectively, and the ratio between them was 2.761. Overall, the pentagon area-based composite score was 310% higher than for the biological data (Table S1).

Likewise, for simulations in which the abundance of the non-motoring crosslinker component, which represented anillin, alone were adjusted (Figure 4B), the contractile rings constricted the dynamic space (30/30) with non-biological kinetics (Figure 4B’). Just as with the simulations with no composition changes, non-motor crosslinker changes alone resulted in a statistically faster normalized maximum constriction speed, of 1.21, which occurred earlier than in the biological or simulation standards, at 25% closure. These simulations also exhibited higher acceleration and deceleration (3.3×10^-2^ %/sec^2^ and 1.1×10^-2^ %/sec^2^ respectively; Figure 4B”, Table S1). The composite score for these was 330% higher than that of the biological standard, comparable to that seen following adjusting tethers alone, and simulations with no composition changes.

Next, we simulated rings in which only the abundance of the actin-like filament changed (Figure 4C). These simulations successfully constricted beyond 90% closure (30/30) and closely recapitulated the constriction kinetics of our biological dataset (Figure 4C’). Overall, the normalized maximum speed was close but statistically distinguishable from the biological and simulation standards at 1.07. Its occurrence at 45% closure, on the other hand, was indistinguishable from both standards (Table S1). Acceleration and deceleration were 1.2×10^-2^ %/sec^2^ and 1.1×10^-2^ %/sec^2^, respectively (Table S1). The ratio between acceleration and deceleration was 1.1, which was statistically different from the biological standard but not the simulation standard (Table S1). The parameter pentagon for simulations in which the abundance of only the actin-like filaments was modulated was only 40% larger than the biological standard and 10% larger than the simulations standard (Figure 4C”; Table S1).

Finally, we simulated contractile rings in which only the motor ensemble abundance was modulated (Figure 4D). 30/30 of these simulations successfully constricted the dynamic space beyond 90% closure. Their contractile kinetics closely matched the kinetics of the biological and simulation standards (Figure 4D’, D”). The normalized maximum speed was statistically different from both standards, at 0.95; maximum speed occurred at 44% closure which was different from the biological standard but not the simulation standard (Table S1). Similarly, acceleration and deceleration were 1.3×10^-2^ %/ sec^2^ and 1.2×10^-2^ %/sec^2^ respectively (Table S1). For these simulations, the ratio between acceleration and deceleration was 1.1 (Table S1). Altogether, the parameter pentagon for these simulations was only 10% larger than the biological data (Figure 4C”; Table S1).

### Omitting abundance dynamics of ring components has distinct effects on constriction

Our data suggested motor and filament abundance dynamics are sufficient to generate more accurate constriction kinetics. We next tested whether they are necessary to generate such accuracy. Since tether and crosslinker abundance dynamics did not influence constriction kinetics in our previous simulations, we set up simulations wherein all components but tethers (Figure 5A) or all components but crosslinkers (Figure 5B) were modulated as a baseline. Omission of the abundance dynamics of either component alone resulted in constriction kinetics similar to the biological data (Figure 5A’, B’), and an overall composite score 30% above the biological dataset, which was indistinguishable from the simulation standard (Figure 5A”, B”; Table S1). Next, we ran simulations where all components were modulated except for filaments (Figure 5C) or motors (Figure 5D). Only simulations with static motor abundance were worse at recapitulating biological kinetics with a 50% increase in composite score (Figure 5D”, D”; Table S1). Altogether these results suggested that the dynamic abundance of motor ensembles specifically is of key importance for realistic constriction kinetics.

**Figure 5.**
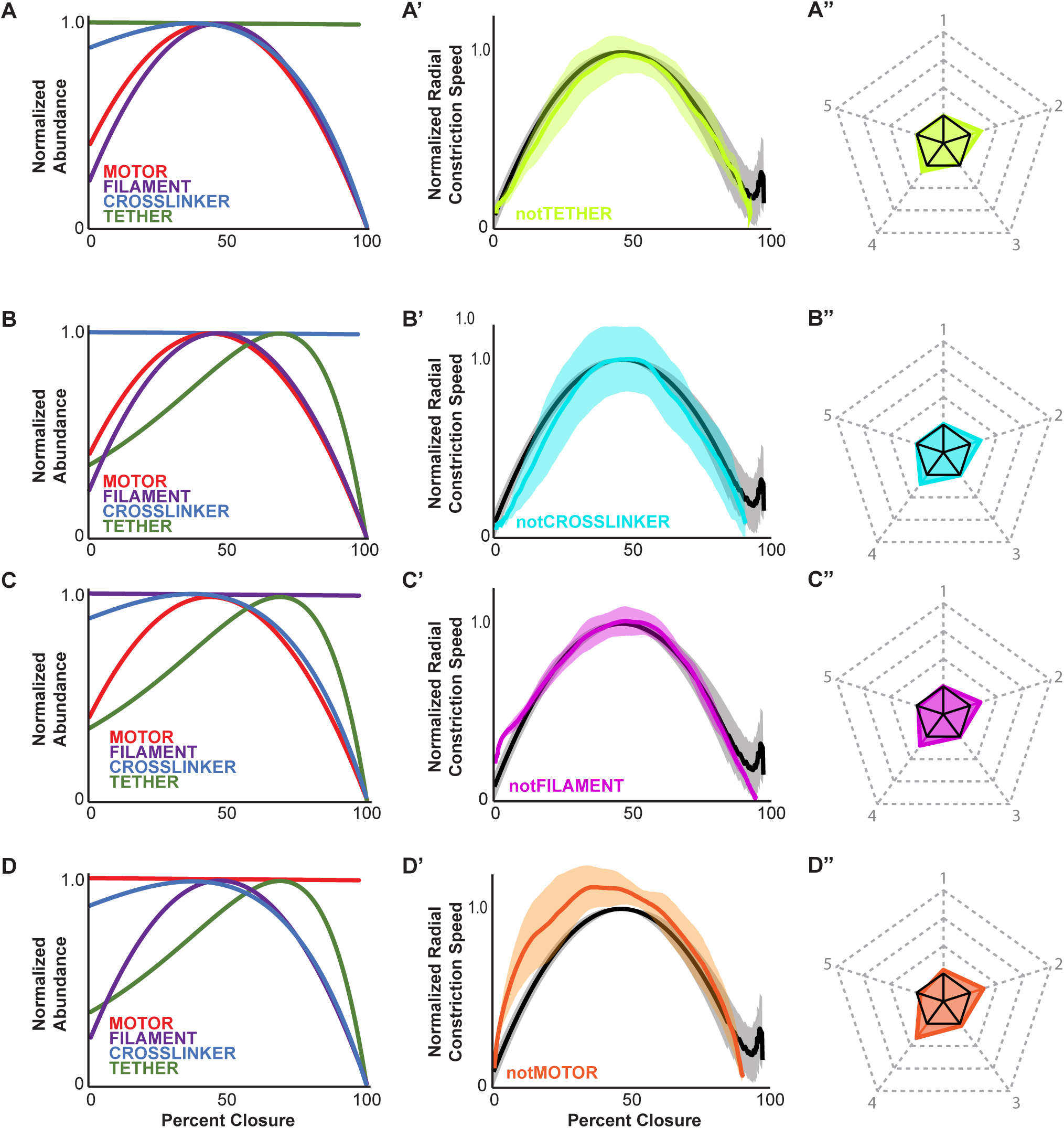
Omitting abundance dynamics of different ring components has distinct effects on constriction. A-D) Normalized component abundance over closure input for simulations with conditions in which dynamics of each of the four main ring components was individually omitted. A) All components dynamic except tethers. B) All components dynamic except crosslinkers. C) All components dynamic except F-actin-like filaments. D) All components dynamic except motor ensembles. A’-D’) Plots of average constriction dynamics curves of simulated divisions (correspondingly colored curves) versus the curve from the biological dataset (black line). Lighter colored areas around solid curves: mean ± standard deviation. A”-D”) Pentagons generated from the five contractile metrics measured from each simulation setup (color-matched) compared to the biological standard metrics (black solid pentagon). Closure dynamics graphs include the curve generated from image data as a standard for comparison (black curve); lighter colored areas represent the mean plus or minus the standard deviation. Solid black pentagons: biological standard. n = 30 for all.

### Motoring activity or fast filament treadmilling is required for realistic constriction kinetics

NMMII motor complexes contribute several activities to actomyosin network kinetics, including crosslinking, bundling and translocating F-actin (Billington et al., 2013; Nagy et al., 2013; Beach and Hammer, 2015; Stam et al., 2015; Melli et al., 2018). All these activities could underlie the importance of implementing myosin component abundance for accurate constriction kinetics. Thus, we next tested whether motoring activity was necessary for this effect by simulating contractile rings bearing myosin-like complexes with no motoring activity. Myosin-like components were unchanged except that their motoring speed was set to 0 nm/s; this ensured that fiber binding dynamics were unaltered but that the myosin-like components did not directly generate force representing the power stroke. These simulations were run with all component composition changes (Figure 6A). Under these conditions, filament networks were contractile, but constricted approximately 16-fold slower than networks in which myosin-like complexes possessed motoring activity (Table S1). Constriction speed remained consistent over time and simulated rings did not close past 45% (20/20) during the simulation time that allowed all other simulated rings to close (Figure 6A). Previous biological and *in silico* work reported contractility in the absence of NMMII (or simulated myosin) translocation. One possible explanation for this observation is that faster F-actin treadmilling can generate force through Brownian ratcheting and persistent attachment of end-bound myosin (Mendes Pinto et al., 2012; Oelz et al., 2015). We tested this possibility by simulating contractile rings with motor-dead myosin-like components as before, and increased filament treadmilling five-fold. Under these conditions, the contractile rings constricted fully (30/30) with kinetics comparable to our biological measurements. The normalized maximum speed was statistically different from both biological and simulation standards, at approximately 1.14, but occurred at point indistinguishable from the simulation standard, at 44% closure (Figure 6B-B”; Table S1). Thus, as in the *in vivo* and other simulation regimes, the absence of motor activity can be compensated for by filament treadmilling and crosslinker end-tracking.

**Figure 6.**
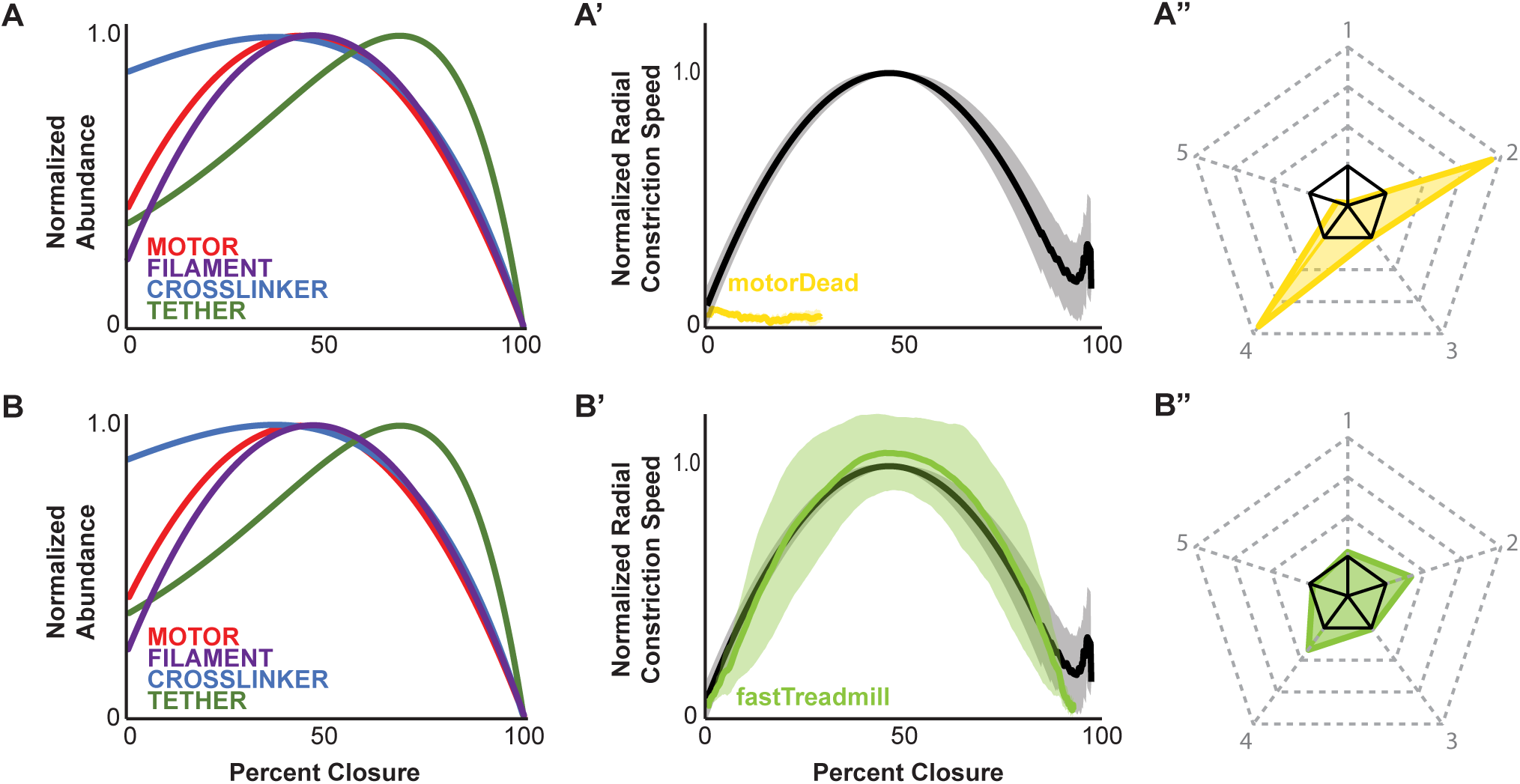
Motor activity or fast filament treadmilling is necessary for normal contraction dynamics *in silico.* A, B) Dynamic abundance of all components was implemented in simulations. A’) Graph of average constriction dynamics of simulations with motor-dead motor ensembles. B’) Graph of average constriction dynamics of simulations with motor-dead motor ensembles and fast-treadmilling F-actin-like filaments. A”, B”) Pentagon showing comparison of five constriction metrics to the biological standard. Closure dynamics graphs include the curve generated from image data as a standard for comparison (black curve); lighter colored areas represent the mean plus or minus the standard deviation. Solid black pentagons: biological standard. n = 30 for all.

### Switching abundance of motors with crosslinkers changes contraction kinetics

While myosin and F-actin abundance increased in a linear fashion unlike anillin and septin, whose abundance changed exponentially (Figure 1B-E), the dynamic density curves of all four components shared the characteristics of accumulating to a single peak during the middle third of ring constriction, followed by loss of components (Figure S3A’-D’). To test the importance of the measured abundance dynamics for each component to *in silico* contractile ring constriction kinetics, we swapped the abundance dynamics of motor components (linear change) with crosslinkers (exponential change; Figure 7A). These simulated rings constricted successfully (30/30) but exhibited drastically altered constriction metrics (Figure 7A’, A”). Normalized maximum constriction speed was 2.1- fold faster than our biological or simulation standards, and was reached at 39% closure, much earlier than either the biological standard or the simulation standard (Table S1). Acceleration and deceleration were both markedly faster than both our biological measurements and the simulations lacking abundance dynamics at 3.0×10^-2^ %/sec^2^ and 2.4×10^-2^ %/sec^2^ respectively (Table S1). In conclusion, myosin abundance dynamics measured in the *C. elegans* zygote were crucial for simulating contractile ring constriction with realistic kinetics.

**Figure 7.**
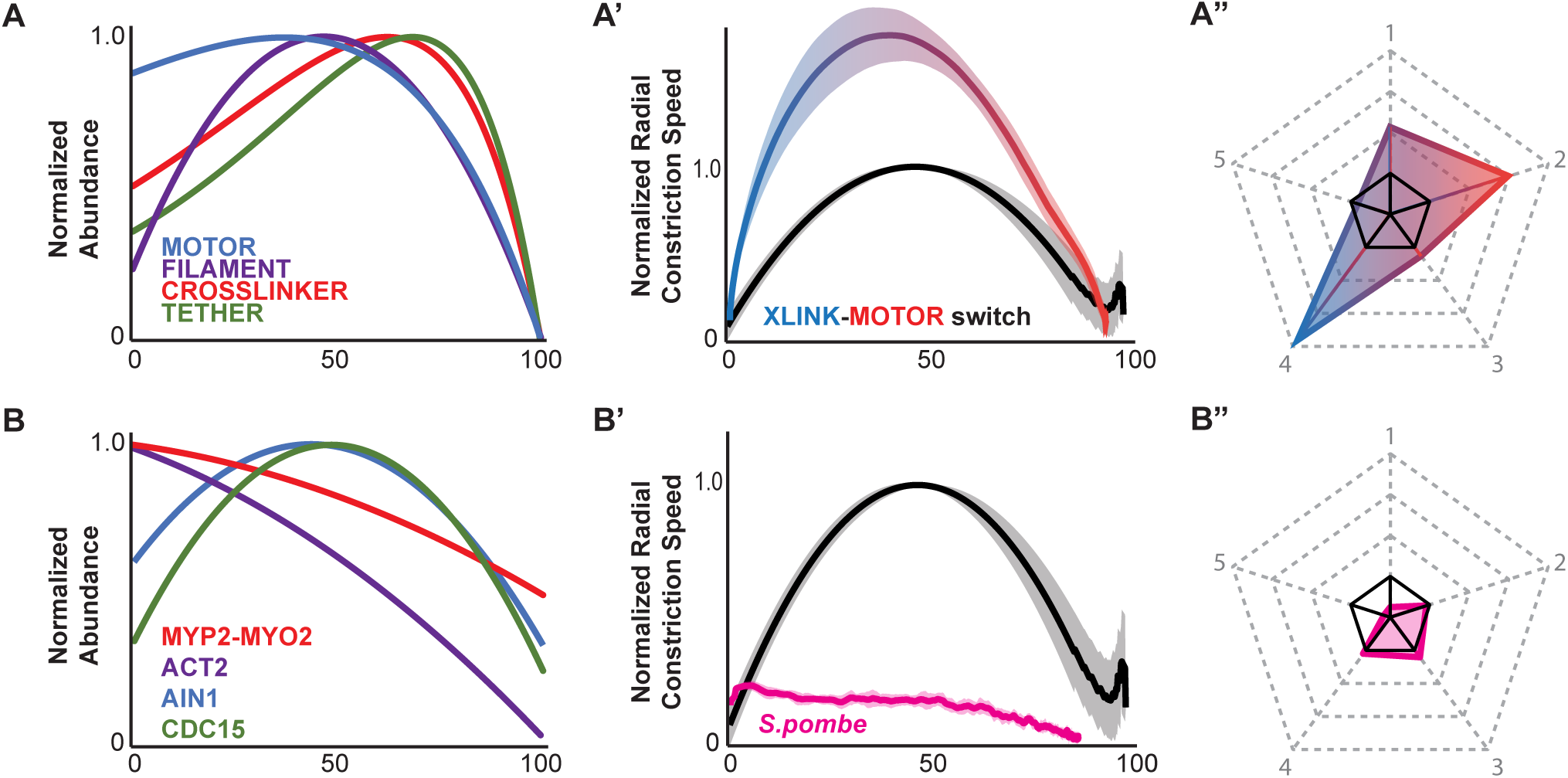
Altered abundance dynamics affect simulated contractile ring contraction. A-A”) Effects of switching motor and crosslinker abundance dynamics in simulations. A) Graph of normalized abundance showing that all components were allowed to change abundance but motors and crosslinkers now have switched their respective curves. A’) Graph of average constriction dynamics of simulations from A. A”) Pentagon showing comparison of five constriction metrics to the biological standard. B-B”) Effects of swapping out C. elegans abundance dynamics with approximations of fission yeast abundance dynamics. B) Graph of normalized abundance showing that all components were allowed to change abundance but with alternate dynamics calculated from fission yeast literature. B’) Graph of average constriction dynamics of simulations from B. B”) Pentagon showing comparison of five constriction metrics to the biological standard. standard. All closure dynamics graphs include the curve generated from image data as a standard for comparison (black curve); lighter colored areas represent the mean plus or minus the standard deviation. Pentagon plots include the biological standard in solid black.

## Discussion

### Ring composition as a predictor of constriction kinetics

Here we report that incorporating changes in global composition into simulations of the contractile ring throughout its constriction, lead to more realistic constriction kinetics. This is noteworthy considering that abundance and constrictions were both measured in the same cell type. Although our simulation parameters were compiled from published studies on a range of cell types (Wu and Pollard, 2005; Guo and Guilford, 2006; Kovacs et al., 2007; Billington et al., 2013; Melli et al., 2018), incorporating the measurements from the *C. elegans* zygote dramatically improved the resemblance between the simulated contraction and the kinetics measured in the same cells. Moreover, by swapping our component dynamics measurements from the *C. elegans* zygote with those made in *S. pombe* (Wu and Pollard, 2005; Courtemanche et al., 2017) we observed constriction kinetics more closely resembling those of fission yeast cytokinesis (Figure 7B-B”). These results highlight the importance of accurately representing the contractile ring composition in any simulation of the actomyosin contractile ring aiming to reproduce the closure dynamics.

### Motors and filaments are the backbone of contractility

The components contributing most prominently to simulated constriction kinetics were those representing F-actin and NMMII, rather than the components emulating tethers and non-motoring crosslinkers. In fact, simulating proper contractile ring composition of either filament or motor alone was sufficient to yield the biological constriction kinetics. The effectiveness of modulation of actin-like filaments likely stems from their significant role as a substrate to which motors and crosslinkers bind and generate forces. Biological f-actin can also generate force directly via treadmilling since some crosslinkers can associate persistently with depolymerizing ends (Melli et al., 2018). Therefore, the abundance of biological and simulated filaments directly and positively relates to network contractility. Similarly, modulation of motor abundance is likely important due to the several contributions of multimeric NMMII motor ensembles to actomyosin network architecture and direct force generation. *In silico,* as well as *in vivo*, contraction speed is substantially affected by changing the number of active motors present on the contractile ring (Descovich et al., 2018; Osorio et al., 2019). Furthermore, because NMMII filaments likely have 10-30 motor pairs, they contribute substantially to network connectivity. We previously showed that discretely simulating the motor domains that comprise ensembles optimized contractile ring constriction, unlike coarse-graining the collection of motor subunits as a single entity (Cortes et al., 2020). Finally, in our simulations, even motor-dead motor complexes increase contractility of treadmilling actin filaments by being strong end-binders (Melli et al., 2018). Here, motor components were simulated with binding and unbinding rates informed by NMMII binding dynamics measurements (Guo and Guilford, 2006; Melli et al., 2018), and with persistent binding of ends such that they consistently track shrinking ends until their internal dynamics dictated unbinding. Therefore, if biological NMMII motors are indeed strong end-binders, we would expect their abundance to have profound effects on constriction kinetics regardless of motoring activity as has been shown with partial global depletions by RNAi (Descovich et al., 2018).

Our *in silico* results with motor-dead motor components highlight two regimes under which biological constriction can occur. First, with slower actin filament treadmilling, motoring activity was necessary to generate sufficient circumferential contraction to drive radial constriction. In this regime, loss of motor activity resulted in very slow contraction and constriction, which were roughly 16-fold slower than constriction with motoring activity unaffected (Figure 5C’, C”). In these simulations, constriction was driven by a Brownian ratchet mechanism where motors bound to depolymerizing filament ends indirectly generated constriction through the active shrinking of the depolymerizing filament end. However, because treadmilling was slow in these simulations, constriction did not reach 30% closure within 300 seconds of simulated time, far slower than *C. elegans* zygotic division. This observation is partly in line with the necessity of motoring as has been reported in *C. elegans* (Osorio et al., 2019). The second regime, where actin treadmilling is far faster and therefore motoring activity becomes dispensable for constriction kinetics, was also explored in our simulation work. With five-fold faster filament treadmilling, simulations with motor-dead motor components were almost indistinguishable from our simulations with motor activity and slower filament treadmilling (Figure 5’D’, D”). Here again, constriction was driven by a Brownian ratchet mechanism, but now with faster treadmilling and therefore faster filament end shortening, constriction was able to generate enough radial force to constrict the deformable dynamic space. Our work suggests that systems where motoring activity is dispensable should have faster actin treadmilling or filament shrinking than systems where motoring activity is necessary for constriction to proceed as normal.

It is worth noting that while modulating levels of both motors and filaments profoundly affected contractile kinetics, modulating either alone was sufficient for recapitulating biological kinetics in our simulations. Furthermore, in simulations where the amounts of all components except for motors or filaments were allowed to change we saw that motors were more necessary for accurate kinetics than filaments. Altogether, these data suggest that the contributions from motor and filament components to constriction are partially redundant, but ring kinetics are slightly more sensitive to motor tuning than filament tuning all other things being equal. This denotes a greater contribution by motors either to force generation than filaments by treadmilling, or to structure and force transmission by crosslinking of filaments. It is possible that with further fine-tuning of the model and the biological data these possibilities could be tested.

Modulation of tethers and non-motor crosslinker composition, on the other hand, had very little effect on overall constriction kinetics. This is unsurprising as these components did not directly contribute to force generation and provided less connectivity, per component, than motors to actin-like filaments (maximum of 2 [crosslinker] or 10 [tether] versus up to 30 [motor]). Similarly, since they are smaller than motors and filaments, non-motor crosslinkers and tethers likely also play less of a role in the deceleration phase of constriction, when network compaction and curvature slow contraction

### Simulation advancements in this work

Simulations of contractile rings have advanced our understanding of contractility. However, whether contractile networks have been represented as active gels (Turlier et al., 2014; Sain et al., 2015) or nematic active gels (Salbreux et al., 2009; Reymann et al., 2016; Dorn et al., 2016), or modeled as discretized agents in ring formation (Vavylonis et al., 2008; Bidone et al., 2017) and constriction (Mendes Pinto et al., 2012; Oelz et al., 2015; Ennomani et al., 2016; Belmonte et al., 2017; Nguyen et al., 2018; Descovich et al., 2018; Cortes et al., 2020), it has been generally assumed that the amount of contractile ring components, and thus global composition of the whole apparatus, remains unchanged throughout the process under study. In addition, contractile networks have largely been modeled independent of their relevant bounding space, such as the plasma membrane, in vivo.

Here, we increased the realism of simulations by incorporating a simplified approximation of membrane deformation energetics derived from the evolving geometry of a capsuloid cell as it constricts at its equator. This derivation, while simplistic and requiring several assumptions, is in general agreement with the kinetics described by the active gel models. We paired this with a deformable space in Cytosim, which can interact with confined filaments and changes in shape in response to forces from filament networks (Dmitrieff et al., 2017). The result was a deformable cylinder space whose resistance to deformation was dependent on the instantaneous size of the space radius (as a proxy to contractile ring radius). Finally, we advanced the field by incorporating adaptive component adjustments into simulations of actomyosin constriction. By pairing the expected amount of network components to the size of the deformable space, we generated simulations that self-regulate, adding or removing some amount of each component type to match user-defined functions dependent on the dimensions of the bounding space. We set these functions to reflect our *in vivo* measurements; future work will involve incorporating measurements from other cell types or biological contexts and help make predictions about specific conditions such as temporal depletions or over expressions.

### Closing remarks

The implications of this work are subject to a few limitations that need addressing in the future. First, while our abundance measurements come directly from *C. elegans*, the biophysical measurements used to construct our *in silico* contractile rings originate from a mixture of animal and fission yeast work. Our ability to reproduce close approximations of *C. elegans* zygotic division constriction kinetics highlights the universality of the mechanisms that give rise to cytokinesis, but measurements of *C. elegans* NMMII and other proteins will strengthen the predictive power of our simulations. Second, we incorporated a simple geometric model to approximate the force that resists constriction of the capsuloid embryo along the cleavage furrow. This model made assumptions about the scale of membrane tension and viscosity of the *C. elegans* zygote, based on reasonable approximations from the available literature (Yoneda and Dan, 1972; Hiramoto, 1975; Daniels et al., 2006). Again, it would be beneficial to measure the cortical stiffness of the *C. elegans* cortex and contractile ring. In addition, while we report the importance of realistically modeling global component dynamics, we did not explore local component abundance or dynamics. Future work will compare simulated ring ultrastructure to local compositional changes in vivo, such as dynamics of contractile units, as have been reported in cells (Capco and Bement, 1991; Wollrab et al., 2016; Henson et al., 2017).

## Methods

### MM1. Generation of microfluidic chip template by photolithography

Microfabrication of microfluidic imaging traps was performed at UNC under the guidance of Dr. Matthew DiSalvo and Dr. Nancy Allbritton. Chamber design was adapted from an embryo trap originally made for high-throughput embryo sorting (Cornaglia et al., 2015), but with altered channel dimensions to trap embryos in a vertical orientation instead of a horizontal orientation (orthogonal versus parallel to the focal plane, respectively; Figure 1A). Modifications were made to trap width and channel size to ensure proper flow for trapping and suction. Once finalized, the design was printed on glass (FrontRange Photomask), which was used as a mask for photolithography (Figure 1A, Figure S1C).

For template creation, a large 50 x 75 mm glass slide was first cleaned thoroughly with 99% isopropanol (IPA) and air-dried using nitrogen gas. The cleaned slide was then placed in a plasma chamber for 3-5 minutes in order to clean the top layer of the glass surface and improve deposition of the photoreactive resin (photoresist 1002F-50; Pai et al., 2007). The glass slide was then placed, and vacuum sealed, onto a minicentrifuge and an aliquot of photoresist approximately 1 inch in diameter was placed at the center of the slide (over the vacuum sealed pedestal). The slide was then spun at 200 rpm for 10 seconds to spread out the bulk and then at 1890 rpm for 30 seconds to create a uniform coating of photoresist 80-100 microns thick.

Following this initial coating, the photoresist hardened while the slide incubated at 95°C for 1 hour, protected from dust or other debris. After incubation, the patterned photomask was carefully placed onto the photoresist and the slide was exposed to 260 mJ UV light (∼300 nm) at 450 mW power. Following exposure, the photomask was carefully removed from the slide and the slide was incubated for an additional 30 minutes at 95°C.

Next, the glass slide was washed on an orbital shaker with fresh resin solvent for 90 seconds and rinsed with used solvent for 30 seconds, ensuring even coverage across the whole side surface. Finally, the slide was washed twice with IPA for 15 seconds and briefly air dried. After washing off uncured photoresist, the slide was examined for proper feature retention with a stereoscope at 10x magnification. If successful, the slide was then incubated overnight at 95°C for 30 minutes then at 120°C overnight to fully cure and harden.

Next, the slide was silanized with trichloro(octyl)silane (Sigma-Aldrich) overnight in a vacuum chamber under vacuum in order to reduce adhesion of polydimethylsiloxane (PDMS; Sylgard 184, Corning) to the finished template. Finished slides were kept in a vacuum chamber for up to 1 month without need for re-silanization.

### MM2. PDMS casting of microfluidic devices

To image *C. elegans* zygotes on the Mizar TILT system (Fadero *et al*., 2018), it was necessary for PDMS microfluidic chips to have enough thickness such that a glass coverslip could be securely affixed to the side of the PDMS, orthogonal to the imaging surface coverslip (the two coverslips being the entry point for the light sheet, and against which the embryo rests, respectively: Figure 1A). Thus, PDMS of the desired thickness, which was about 10 mm, was cast in an aluminum chamber within which we could place a template, lithograph side up. Proper TILT imaging with minimal scattering of the illumination light sheet requires that the light sheet enter the sample chamber roughly parallel to the imaging plane, through a nearly orthogonal, flat, and flawless surface. We therefore designed the casting chamber to have mirror-finished interior surfaces with a slight tilt (2.3° offset from orthogonal); this resulted in a nearly flawless lateral PDMS surface onto which a coverslip was later affixed. For ease of microfluidic chip removal, the casting chamber was made fully disassemble-able.

For casting, the back surface of a glass template was lightly coated in vacuum grease and carefully centered onto the bottom of the casting chamber bottom panel. The casting chamber was then assembled with care to ensure that the template slide did not shift. Uncured PDMS was then poured slowly into the chamber to the desired height and degassed under vacuum for 3-5 minutes. After degassing, the PDMS was cured in an incubator for 2 hours at 90°C. Our templates were constructed to produce six microfluidic chips per casting; 10 minutes prior to removing the cast PDMS from incubation, six 60 x 25 mm coverslips along with six 22 x 22 mm coverslips (all #2) were prepared for binding to the PDMS. All coverslips were carefully washed with 100% ethanol and allowed to air dry. Dry coverslips were placed in a plasma chamber for 3 minutes for surface cleaning, all coverslips were then placed, same side up, into an airtight container for temporary storage.

All steps involving preparation of hardened PDMS into fully finished microfluidic chips with coverslips were performed in a laminar flow hood to avoid contamination with dust. The PDMS casting was removed from the incubator and the casting chamber was carefully disassembled. The hardened PDMS was then carefully removed from the template and care was taken to avoid contact between dust or debris and the bottom or side PDMS surfaces, which were nearly flawless mirror-finished, and to which coverslips would be affixed. The six microfluidic chips were then cut apart using a new disposable scalpel. A 1.5 mm biopsy punch was used to create the inlet and outlet channels from the PDMS. Finally, a 22 x 22 mm coverslip was carefully placed, plasma-cleaned side towards the PDMS, onto the angled lateral surface of each chip, and a 60 x 25 mm coverslip was similarly placed onto the bottom surface of the PDMS with the PDMS chip roughly centered on the slide. The casting chamber created a small lip of PDMS because the bottom surface of the chamber is roughly 1.5 mm larger than the template glass slide on all sides. Thus, each PDMS chip had a lip of 1.5 mm width by ∼1 mm thickness which allowed for proper placement of the bottom coverslip without impinging on the vertical coverslip. Once both coverslips were placed on all microfluidic chips, assembled chips were carefully placed at 60°C for 3 hours to cement coverslips to the PDMS. Completed devices were stored indefinitely in airtight containers, protected from dust before and after being used for imaging, and were reused indefinitely with proper storage and maintenance, flushing with distilled water after every use.

### MM3. C. elegans strains and maintenance

The following *C. elegans* strains were used for the live cell imaging:

**SWG007:** (nmy-2(cp8 [nmy-2::GFP unc-119+]) I; gesIs001 [Lifeact :: mKate2]) (Reymann et al., 2016)

**NK2228:** (unc-59(qy88[unc-59::GFP-C1::3xflag::AID+loxP])) (Chen et al., 2019)

**MDX87:** (unc-59(qy88[unc-59::GFP-C1::3xflag::AID+loxP]); Si57[pEZ152; pani-1:mkate2::ANI-1; cb-unc-119(+)]IV). This strain resulted from a cross of NK2228 with ZAN103.

**MDX29:** (ani-1(mon7[mNeonGreen^3xFlag::ani-1]) III)

**MDX82:** (unc-119(ed3) III; ltIs81 [Ppie-1::gfp-TEV-Stag::ani-2; unc-119 (+)] ;gesIs001 [Lifeact :: mKate2]; mgSi3[tb-unc-119(+) pie-1>gfp:utrophin] II.; unc-119 (+)] III; ltIs37 [pAA64; pie-1/mCHERRY::his-58; unc-119 (+)] IV). This strain was made by crossing SWG001 (Reymann et al., 2016) with JCC719 (Jordan et al., 2016).

*C. elegans* strains were maintained as previously described (Munro et al., 2004).

### MM4. Dissection and embryo staging

L4 hermaphrodites were grown overnight at 25°C. The following day, 2-4 gravid adults were placed on a watch glass with autoclave-sterilized M9 buffer [22 mM KH2PO4, 42 mM Na2HPO4, 85 mM NaCl, 1 mM MgSO4]. Following an initial cleaning in the M9, worms were dissected with 20-gauge needles and eggs were pooled at the center of the watch glass. Using a hand-drawn fine capillary tube and mouth aspirator, zygotes in which pronuclei had met or which were in pseudocleavage as identified with a stereoscope were aspirated and placed directly into the inlet channel of a microfabricated PDMS imaging chip. A 20 mL syringe full of M9 was then attached to the inlet channel via microfluidic tubing and a flat head needle. A syringe-controlled flow of M9 through the microfluidic chip for zygote trapping and provided a liquid well to maintain fluid pressure and ensure zygotes stayed trapped in the imaging channels. A small volume of M9 was then slowly pushed through the microfluidic chip while observing through a dissection microscope to check for flow-through of worm zygotes. If zygotes were observed to be caught in an imaging cup, flow was immediately stopped; if zygotes flowed through without becoming trapped, flow was reversed and flow-through was repeated more slowly. Zygotes trapped in an angled or horizontal position (long axis horizontal) were coaxed to stand up vertically (long axis vertical) via quick short pulses of forward and reverse flow. After trapping zygotes in a vertical orientation, the microfluidic imaging chamber and attached syringe were carefully placed on the stage of an inverted microscope for time lapse imaging.

### MM5. Imaging on Mizar TILT

Microfluidic chips were placed directly onto a Nikon Ti Eclipse inverted microscope (Nikon Instruments) with a Mizar TILT modular light sheet (Mizar; Fadero et al., 2018). A 10x air objective (Nikon Instruments) was used to locate zygotes in the imaging chip; a 60x planApo oil immersion objective (Nikon Instruments) with an N.A of 1.42 was used for acquisition along with a Prime 95B CMOS camera (Photometrics). Micromanager was used to operate the microscope body, shuttering system (Sutter), and light sheet box. Modular power lasers (OBIS) at 488 nm and 594 nm wavelengths were used for imaging; zygotes were located via brightfield. Zygotes were scanned in the Z-dimension to check for pronucleus (or metaphase plate) location in order to verify correct staging and orientation. Zygotes were only imaged and analyzed if their contractile ring formed close to on-plane or on-plane with the imaging light sheet. Once a properly aligned zygote was identified, it was observed via light-sheet in order to approximate the range in Z over which the contractile ring was likely to form based on the absence of cytoplasmic signal at the location of the pronuclei or based on enrichment of the cortical fluorescent signal (Figure 1A, B). Immediately upon detection of the contractile ring, a time lapse with Z-stack acquisition (11-25 optical sections, 200 nm apart) was initiated. Imaging was continuous with 150 millisecond exposures, resulting in timesteps of approximately 3-7 seconds. We used an OBIS laser module with a 150mW 561nm laser at 40% power for red flours and a 150mW 488nm laser at 25% power for green flours.

### MM6. Image pre-processing

All image pre-processing was performed in FIJI (ImageJ). Image sequences were trimmed in x, y, z, and time to contain only the contractile ring. The resultant 4D images (XYZT) were then Z-projected (FIJI>Image>Stacks>Z Project) for maximum signal intensity in the Z-dimension for all time points, resulting in a 3D stack which contained 2D projections for each time point (XYT).

For segmentation of contractile ring signal, we also generated a minimum intensity projection from the original 4D stack (XYZT). The resulting 3D stack was minimum intensity projected again, to generate a 2D image of the minimum signal for each XY pixel over all Z and T dimensions. This 2D image was then subtracted from each time point in the maximum intensity projection time series. The “result” stack (XYT; maximum intensity minus minimum-minimum intensity) and the original maximum intensity projection stack (XYT) were saved for processing through a custom pipeline to segment contractile ring signal, report signal intensity and calculate contractile ring kinetics.

### MM7. Contractile ring measurements

Custom code was adapted to segment the contractile ring in end-on orientation throughout cytokinesis (Ishikawa and Geiger, 1998; Kang Li et al., 2006). Processed stacks were segmented by a neural network that fits a polygon to the shape of the contractile ring and digitizes the resulting shapes into a binary stack with the same pixel and temporal dimensions as the input. These ring stacks are then exported as TIFFs and further processed in FIJI (Figure S1D, third panel).

Ring stacks were first dilated (FIJI>Process>Binary>Dilate) in order to thicken the ring masks to be 6-8 pixels thick (Figure S1D, 4^th^ panel). For each frame of the maximum intensity projection stack, the corresponding frame of the mask was used to select the ring region from where average pixel intensity, Feret’s diameter, area, and perimeter were measured. For each maximum intensity projection stack, the average pixel intensity of the background was also measured in a 10×10 pixel region of interest in the cytoplasmic signal away from the contractile ring. The background measurement was taken for each time point in the series and subtracted from the average contractile ring pixel intensity for the corresponding time point.

The relative amounts of several contractile ring components throughout cytokinesis were quantified as a function of percent closure with respect to initial cell radius (Figure S1A, B)

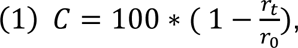

where r_t_ is the radius (or ½ of the ring diameter; herein the diameter along the major axis of the mask shape) of the contractile ring at the current time point and r_0_ is the initial radius estimate (Figure S1B). Component fluorescence intensity for a given cell was normalized to its value at 50% ring closure.

Biological data from all imaged strains were pooled to generate our *in vivo* contraction dynamics estimates. First, each individual cell’s speed curve was generated from a second-degree polynomial fit of its speed over percent closure. Each curve was then normalized to its own internal maximum speed (such that all curves reached a peak speed of 1.0). All cell data were then averaged to generate a mean fit of speed over percent closure for all points between 0% (fully open) and 100% (fully constricted). For each averaged speed, the standard deviation was calculated and the mean speed plus/minus the standard deviation was reported.

### MM8. Calculating relative fold change from normalized intensity curves

To generate population average normalized fluorescence intensity curves for each fluorescent marker, a linear regression was fit to the normalized intensity data for actin and myosin curves; the log of the normalized intensity data was fit for septin and anillin curves. The fitted lines for all cells from each strain were then averaged together to generate the average linear regression of the entire population of data points each for myosin, actin, septin, and anillin. Finally, for septin and anillin, linear fits were exponentiated to derive the average exponential fit.

Population-averaged fluorescence intensity curves served as proxies for the estimated density of each contractile ring component during constriction. We assumed that the contractile ring width stayed roughly constant after constriction begins; therefore, we multiplied the estimated normalized density of ring components by the perimeter of the contractile ring. This served as an estimate of the contractile ring area and provided an estimated amount of each ring component for each timepoint throughout constriction. The resulting normalized density curves relate to the relative, not absolute, change in protein amount for each contractile ring component (Figure S3). These final curves were used to estimate component amounts in our contractile ring simulations as discussed in MM10-2.

### MM9. Base model description

Our agent-based models of cytokinesis were built within Cytosim (www.cytosim.org), an Open-Source stochastics physics engine used to simulate cytoskeletal networks. The base Cytosim code can be found on Gitlab (https://gitlab.com/f-nedelec/cytosim). Significant modifications were however needed to run our specific simulations (discussed in the next section), for which code is available on Gitlab (https://gitlab.com/dbcortes/cytosimDev-6_21/-/tree/paper2022).

Contractile ring simulations were run in 3D (discussed below) and included agents representing the main molecular components of animal cytokinetic rings (Table S2) interacting with an outer cylinder of variable radius, representing the plasma membrane of a constricting cell.

F-actin was simulated as a treadmilling filament initially 500 ± 200 nm in length, exhibiting stable treadmilling where the plus (barbed) end assembly and minus (pointed) end disassembly both occur at 10 nm/s. Septin, a crosslinker and membrane tether in the contractile ring, was simulated as a 100 nm long filament with ten evenly spaced actin-binding components (herein, binding domains are referred to as “couples”). Anillin, a scaffold protein, was simulated as a 40 nm long filament with two actin-binding couples and one motor-binding couple affixed on the filament backbone. A final simple crosslinker was simulated as a short 20 nm-long filament bearing two actin-binding couples. Non-muscle myosin II (NMMII), the crosslinking motor implicated in driving cytokinesis, was simulated as a bipolar filament ensemble (Cortes et al., 2020) as characterized through electron microscopy (Nagy et al., 2013; Billington et al., 2013; Beach and Hammer, 2015; Melli et al., 2018). Each ensemble consisted of 30 motors, with fifteen motors bound to either end of a single 300 nm-long filament. Each motor binds and unbinds actin-like filaments independently as force is exerted across the spring-like linker. Motors had a maximum unloaded speed of 150 nm/s, and lower speeds under load as demonstrated *in vitro* and *in vivo* (Stam et al., 2015; Guo and Guilford, 2006; Billington et al., 2013; Nagy et al., 2013; Melli et al., 2018).

The potential effects of component crowding in 3D were depicted with steric interactions between the filamentous representations of all included ring components. The steric range was 10 nm for all filaments, to match the diameter of actin filaments; thus, component segments within 10 nm sterically repelled one another (Nedelec and Foethke, 2007).

Simple component turnover, in which every 1-2 seconds a single agent of each component type was removed from the simulation space and one new agent was added somewhere in the contractile network at random, was necessary to avoid rupture and collapse of simulated rings (Figure S5A-D), as seen for a membrane-anchored model of the fission yeast cytokinetic ring (Nguyen et al., 2018).

The simplest 3D simulated contractile ring was constructed as a ring-shaped meshwork of isotropic filaments and filament-binders confined within a static viscous space. The simulated contractile ring was 10-fold smaller in cross-sectional area than the *C. elegans* zygote to decrease computation load and time. The simplest simulations estimated functional amounts for each of four ring components (filaments, motor ensembles, crosslinkers, and tethers) based on measured maximum protein abundance in the fission yeast cytokinetic ring (Wu and Pollard, 2005).

### MM10. Cytosim agent-based software modifications

A sample configuration file to run a simulation with the modifications detailed below can be found within our Gitlab repository, titled “example_config.cym.” Modifications made to the Cytosim base code for this work are as follows:

1. A physically realistic method for calculating motor unbinding based on force was previously coded and tested in Cytosim (Cortes et al., 2020). Specifically, fully discretized NMMII motor subunits exhibited catch-slip unbinding dynamics and comprised ensembles of 15 motors on either side of a 300 nm-long semi-rigid filament.
2. Default Cytosim allows users to set event codes to add or remove specific amounts of any component in a time-dependent manner. However, we needed to change to component abundance as a function of contractile ring size, and not time. Thus, an algorithm was added to automatically update the amounts of contractile ring components based on the radius of the deformable bounding space. Component fold-changes (MM10) were converted to component count by estimating a maximum amount for each component based on measurements from fission yeast (Wu and Pollard, 2005; Courtemanche et al., 2017). At every time step (1 millisecond) Cytosim calculated the total amount of each component from the radius of the virtual cell given the normalized fold-change functions known for each component. A user-defined ‘adjust’ function (Supplemental File 1) added or removed members of that component class at random in the whole user-defined space, for removal, or in a specified sub-region, for addition, until the expected amount calculated by the fold-change function was reached. In practice, the adjustment function used a Gillespie decay counter to add some stochasticity to these adjustments, to mimic at least some level of cellular variability.
3. The cylindrical bounding space representing the plasma membrane was dynamic such that it could change dimensions (radius) but not shape throughout the simulation. Dimension change was driven by the forces generated by the associated network, similarly to a previous study (Dmitrieff et al., 2017). In short, the rate of radius change was set proportionally to the force exerted. A three-dimensional finite cylinder with radial symmetry about the Z-axis (herein referred to as ’the cell’) consisted of two circular faces parallel to the XY-plane, defined by a radius R and separated along the Z-axis by a length L, and one cylindrical surface, of matching length, and circumference 2𝜋𝜋R. Simulated components were contained within the circular faces. Along the cylindrical surface, however, components were not rigidly contained and were instead confined to be either within or on the surface through an energy potential, characterized by a scalar stiffness parameter. Thus, components slightly outside the cylinder and internalized surface-confined components exerted forces on the cylinder. Components added to the system were set only near the cylindrical surface, and inside the cylinder. Forces were transmitted to the cylinder by cytoskeleton-like components through their interactions defined by the potential associated with the cylindrical surface. Applying the force potential to actin-like filaments and septin-like tethers was sufficient to tether the entire contractile network tightly to the deformable surface, because these components are connected to the rest of the network via motors and non-motor crosslinkers. For any component point subjected to the confinement force potential, its radial displacement was used to calculate a resultant radial force. This force could be directed either inward or outward (contractile or expansile; Figure S2C). Septin-like tethers were confined on the space surface along all of their points, allowing them to exert both inward and outward force on the space. Actin-like filaments were confined in two ways. First, their plus ends were confined to the space surface, also allowing them to exert either inward or outward force on the space. Second, all other actin-like filament vertices were confined inside the surface, allowing them to exert only outward forces. All other components were only confined inside, thus allowing them to contribute only outward forces as a result of either expansile motion in the contractile ring or crowding of components. Forces acting on the cell’s cylindrical surface were calculated for every confined vertex. At every time point, the inward or outward offset of each vertex was calculated by first projecting to the closest point onto the surface. The offset, *x*, was calculated as the difference between the vertex and its projection (Figure S2C), resulting in a vector that is in a radial direction on the XY plane. The associated force is Hookean with zero resting length:

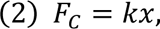

given the confinement stiffness, *k.* The total radial force, 𝐹_𝑅_, is the sum of the radial component of all forces calculated for all confined vertices in the simulated system. The radial component is a scalar that has the same magnitude as 𝐹_𝐶_, positive for outward directed force and negative for inward directed force.

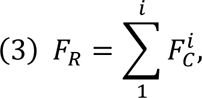
4. To establish the equations governing the radius 𝑅, mathematical estimations of the forces involved in membrane deformation during cytokinesis (Sain et al., 2015) were adapted to Cytosim, creating a custom simulation space class where the radius of the cylinder could change under tension. Using simple geometric descriptors of whole-cell shape as division progresses, and assuming constant total volume, the surface area change of the cell through constriction, as a function of 𝑅, was calculated. Assuming that the energy involved in surface deformation is affected by surface tension, 𝑇, the energy of deformation, 𝐸, of the cell at any given radius was calculated as:

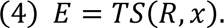 The force of membrane deformation could then be estimated as:

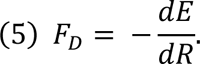

The resultant deformation force was added to the force exerted on the space by contractile ring components,

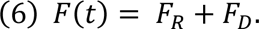

The change in radius, 𝑑𝑅, for each time point was then calculated by multiplying 𝐹(𝑡) by a constant, 𝛼, representing the dampening factors mentioned above:

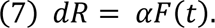

An α value was chosen such that the overall time for simulation constriction scaled with biological data. Finally, the change in radius was added to the current radius at the end of each time step, 𝜏, to calculate a new radius, 𝑅_𝑡+𝜏_, for the following time point:

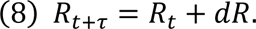
5. The adjust function described in section MM10.2 above could work by incorporating new components into the simulation right under the circumferential surface of the cell. However, because this cell must be wide to simulate the initially wide contractile ring, seeding new components anywhere along its surface resulted in some components never being incorporated into the active contractile ring. We therefore restricted nucleation to a band of width of 2 µm, and thickness of 700 nm. This “equatorial” seeding better reproduced the incorporation of components into the mature contractile ring.

### MM11. Measuring contraction kinetics of simulations

The ‘report’ functionality of Cytosim generates numerical data from saved simulations. A new addition to this is the reporting out of the radius of the dynamic cylinder space into a comma-separated (CSV) format. Our simulations report out the radius every 10 milliseconds and this temporal resolution is much higher than that achieved via live cell imaging of ring constriction (Maddox et al., 2007; Carvalho et al., 2009; Descovich et al., 2018; Rehain-Bell et al., 2017). The reported radius was used to calculate the percent closure (MM7 for description) of the space and the relative speed of constriction at each time point. Speed data were smoothed using a moving mean method with a sliding window of 30 frames (3 seconds), similar to how biological data were treated (Descovich et al., 2018).

### MM12. Estimation of component coverage in dynamic cylinder space simulations

Contractile ring coverage and distribution of motor agents was estimated from color images generated with a hundred millisecond time interval. These images represented the cell as seen from the Z-axis, where it appeared as a circle (Figure S5C, D), akin to a max intensity projection. The images were imported into FIJI (ImageJ) as a time stack and the RGB channels were split into three 16-bit grayscale stacks: one corresponding to the motors, and the other two representing other components which were ignored for this analysis. The motor image stacks were down-sampled to 8-bit grayscale stacks which were then segmented by thresholding to create binary masks for all time points using the Default thresholding algorithm in FIJI (FIJI>Process>Binary>Make Binary). Mask stacks were thickened slightly to fill small holes using binary erosion (FIJI>Process>Binary>Erode). Using the Points from Mask (FIJI>Edit>Selection>Points from Mask) and Fit Circle (FIJI>Edit>Selection>Fit Circle) functions, best-fit circles were generated for all time points and thickened into 1-pixel thick bands using the Make Band functionality (FIJI>Edit>Selection>Make Band). The resultant band regions of interest (ROIs) were then used to measure the mean pixel intensity from the motor mask stack. If the distribution of motor agents along the simulated ring space were 100%, the ROI generated from the circle fitting would be expected to read all pixel values as 255 (max for an 8-bit image). Therefore, the standard for 100% coverage would be a mean pixel intensity of 255. Actual distribution was thus calculated as:

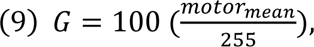

where 𝑚𝑜𝑡𝑜𝑟_𝑚𝑒𝑎𝑛_ is the mean pixel intensity of the motor component image ROI. A visual representation of this same analysis was generated by projecting the two-dimensional view of each individual component type over time. Using this temporal projection, large gaps were made apparent in simulations with inconsistent distribution of contractile ring components.

### MM13. 5-dimensional metric analysis

To quantify the ring contraction kinetics, we defined five metrics: maximum speed, inflection point (percent closure at which maximum speed occurs), maximum acceleration, maximum deceleration, and the ratio between acceleration and deceleration (as a metric of the skew of the kinetics curve; Figure S1E). Measurements were normalized to the biological dataset. Speed had previously been normalized for each individual biological sample to its own internal maximum, so the average maximum speed for 23 measured cells was 1.0 with a standard deviation of zero. All other metrics were measured from the individual standardized speed curve for each cell and the average is reported along with its standard deviation. Normalized metrics were simultaneously displayed in pentagonal multidimensional plots.

Because they were normalized with respect to the biological sample, a simulation that perfectly recapitulated the biology would be represented by a regular pentagon (all radii are 1.0). Maximum speed served as the scaling metric between biological data and the simulation data. The gradient of the smoothed speed curve was taken to calculate the peak acceleration and peak deceleration of each simulation. The maximum speed and inflection point (closure percentage where peak speed was reached) were also tabulated for each simulation. For each simulated configuration, 30 simulations were run to calculate mean and standard deviation data for each of the above-mentioned metrics. For speed and acceleration/deceleration, data from simulations in which all contractile ring component abundances were modulated were set to 1.0 and the scaling factor between its true maximum speed (∼0.054 microns/second) and 1.0 was used to scale speed and acceleration/deceleration from other simulation conditions. Inflection point, which reported the % closure at which maximum speed occurred, was not affected by scaling of the y values of the curve, nor was the slope between acceleration and deceleration since both are scaled by the same factor.

We then generated the pentagons as:

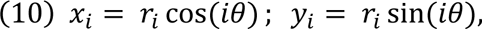

with 𝑖 ∈ {0. . .4} and θ = 2𝜋/5 (Figure S1D). The area of the pentagon was then calculated by breaking it down into five triangles with individual areas:

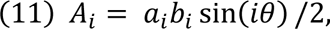

Where 𝑎_𝑖_ and 𝑏_𝑖_ are the lengths of the sides of the triangles (Figure S1F). The area of the pentagon shape is then:

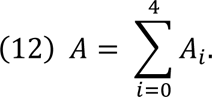

All simulation datasets were analyzed to yield the five metrics from the smoothed speed curve of each individual simulation. These metrics were then averaged for each dataset; thus, for the simulation, standard deviations were reported for all five metrics. The vertices for each simulated dataset pentagon were calculated as before, using the ratio between the simulated condition and the biological data as *r* (Figure S1F). For each comparison, the simulation pentagon was plotted over the biological standard. The 𝚫*r* for each metric was reported to show the fold-increase (or decrease when negative) of each metric. The 𝚫A was reported to show the total relative difference of each simulated dataset from the biological standard. Very small 𝚫A suggested simulations very similar to the biological data, while large 𝚫A suggested simulations very dissimilar from the biological data.

## Supplemental Material

Supplemental material includes a table (Table S1) with the constriction kinetics measurements taken from biological and simulation datasets. We also provide five supplemental figures with further details of our imaging setup, quantification of constriction kinetics for *in vivo* and *in silico* data, and further description of our simulation setup.

## Acknowledgements

The authors thank Dr. Paul Maddox for the innovation of the Mizar TILT system. The microfluidic chips used in this work were manufactured by the researchers with guidance and direct assistance from Dr. Matthew DiSalvo, a previous member of Dr. Nancy Allbritton’s lab. The authors thank the members of the Maddox labs, especially Dr. Michael Werner and Dr. Jenna Perry for their critical evaluation of the research in this work. The automated tracking of contractile rings was based on code implemented by Dr. Florian Jug of the Human Technopole in Italy. Simulation work was made possible through use of the UNC Research Computing cluster, which is supported and maintained by the Research Computing group at UNC.

This work was supported by the NIH/NIGMS R01-102390, NSF 1616661, and the UNC Lineberger Comprehensive Cancer Center through the ITCMS NIH T32 training grant number T32CA009156.

Dr. Amy Maddox declares her relationship with Dr. Paul Maddox, who is Chairman of the Board of Directors of Mizar Imaging, LLC. No other authors declare any conflict of interest.

## Author Contributions

A.S.M was responsible for funding acquisition, supervision, and project administration. D.B.C and A.S.M together, conceptualized the research. D.B.C performed all aspects of the research including the data acquisition, methodology development, software modification, data curation, data validation, and visualization. F.N. provided supervision and validation for *in silico* work performed with Cytosim. D.B.C, A.S.M, and F.N together reviewed and edited the manuscript for content and composition.

---

**Movie 1. Timelapse of NMMII on the contractile ring.** Maximum intensity projection of a 4-dimensional stack showing NMMII (NMY-2::GFP) on the zygotic contractile ring during constriction. Scale bar (bottom right) = 15 μm. Imaging was performed using a microfluidic imaging chip on a Mizar TILT, collecting a full stack every ∼3 seconds. Movie shows 12 frames per second. Related to Figure 1B.

**Movie 2. Timelapse of F-actin reporter on the contractile ring.** Maximum intensity projection of a 4-dimensional stack showing an F-actin reporter (mKate::LifeAct) on the zygotic contractile ring during constriction. Scale bar (bottom right) = 15 μm. Imaging was performed using a microfluidic imaging chip on a Mizar TILT, collecting a full stack every ∼3.5 seconds. Movie shows 10 frames per second. Related to Figure 1C.

**Movie 3. Timelapse of anillin on the contractile ring.** Maximum intensity projection of a 4-dimensional stack showing anillin (mNeonGreen::ANI-1) on the zygotic contractile ring during constriction. Scale bar (bottom right) = 15 μm. Imaging was performed using a microfluidic imaging chip on a Mizar TILT, collecting a full stack every ∼4 seconds. Movie shows 9 frames per second. Related to Figure 1D.

**Movie 4. Timelapse of septin on the contractile ring.** Maximum intensity projection of a 4-dimensional stack showing septin (UNC-59::GFP) on the zygotic contractile ring during constriction. Scale bar (bottom right) = 15 μm. Imaging was performed using a microfluidic imaging chip on a Mizar TILT, collecting a full stack every ∼3 seconds. Movie shows 12 frames per second. Related to Figure 1E.

**Movie 5. Rotating view of agent-based simulation of dynamic component abundance on the contractile ring.** Movie shows a representative simulation of the contractile ring where all four main ring components, motors (red), filaments (purple), tethers (green), and crosslinkers (blue; legend on bottom right) have modulated abundance. Rotation at 0% and ∼25% closure shows how components can be visualized in 3D. The dynamic deformable space is shown in blue. Related to Figure 2.

**Movie 6. Orthogonal view of agent-based simulation of dynamic component abundance on the contractile ring.** Movie shows a representative simulation of the contractile ring where all four main ring components, motors (red), filaments (purple), tethers (green), and crosslinkers (blue; legend on bottom right) have modulated abundance. The dynamic deformable space is not visible from this perspective. Related to Figure 2.

**Movie 7. Orthogonal view of agent-based simulation of static component abundance on the contractile ring.** Movie shows a representative simulation of the contractile ring where all four main ring components, motors (red), filaments (purple), tethers (green), and crosslinkers (blue; legend on bottom right) have preset static abundance. The dynamic deformable space is not visible from this perspective. Related to Figure 3.

**Figure S1.**
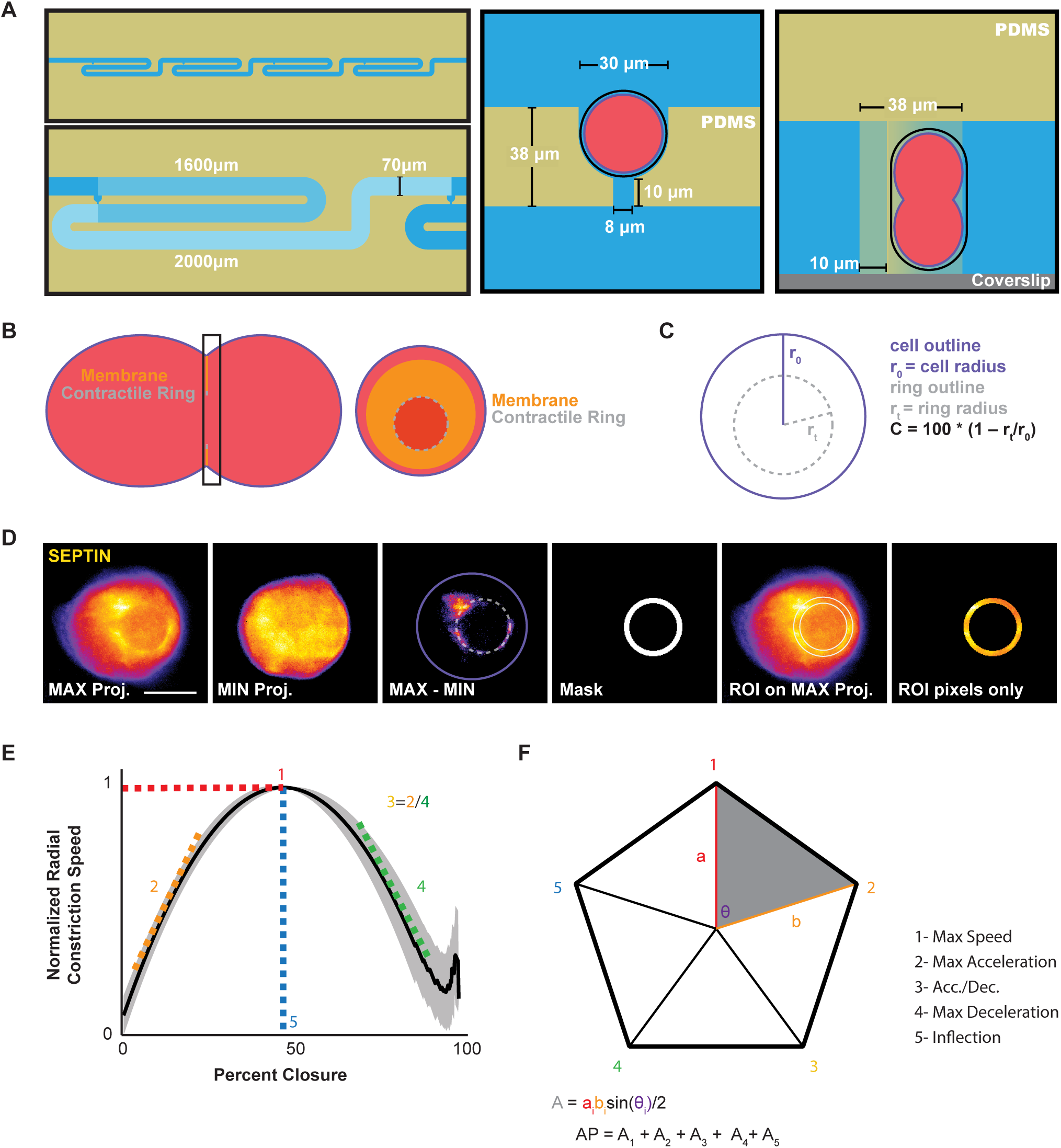
Measurement of contraction dynamics from live cell imaging. A) Zoomed in diagrams of the microfluidic imaging device. Top left: Microfluidic imaging chip with four imaging traps in series. Bottom left: Zoomed in to a single imaging trap with its dimension. Flow channel width is 70 μm, the channel length is 2000 μm before the imaging trap (lightest blue) and 1600 μm after the trap (light blue). Center: Schematic in top-down view of an embryo within the imaging trap. Trap is 30 μm wide within a 38-μm wide column of PDMS. The trap has an outlet channel that is 8 microns wide and 10 μm long, which generates suction through flow. Right: Schematic of side-long view of an embryo within the imaging trap. Trap is approximately 60 μm tall, allowing an embryo of approximately 50 μm in length to stand upright. B) Left: Representative schematic of a dividing *C. elegans* zygote along its long axis. Orange line represents the folded-over membrane along the cleavage furrow. The white tip is the membrane leading edge and contractile ring. Black rectangle: the imaging plane. Right: Projection of the image stack visible from the imaging volume taken along the black rectangle. In this orientation the contractile ring would be visible as a ring (white dotted circle) and the folded membrane representing the cleavage furrow would be the trailing section of fluorescent signal (orange anulus). C) Schematic of measurements taken to calculate closure dynamics. The cell outline is used to calculate r_0_, the initial ring radius. The outline of the contractile ring (gray dotted line) is used to calculate the current ring radius, rt. Percent closure, C, is calculated as a percentage ratio of 1-r_t_/r_0_. D) A single timepoint from a representative timelapse series of a dividing *C. elegans* zygote expressing fluorescently-tagged UNC-59 septin. In order from left to right: Raw image; minimum intensity projection for background subtraction; background subtracted image showing the contractile ring and a fit polygon representing the ring (dotted line); a thickened band mask generated from dilation of the fit polygon; mask overlayed onto the original raw image as an ROI; resulting isolated contractile ring pixel data for measurement of mean pixel intensity and closure dynamics. E) Five metrics of constriction dynamics curves; 1- the maximum speed, 2- acceleration, 3- ratio between acceleration and deceleration, 4- deceleration, 5- the closure % at which maximum speed occurs. F) Standardized pentagon generated from the five contractile metrics measured from biological data set. Scale bar = 15 μm.

**Figure S2.**
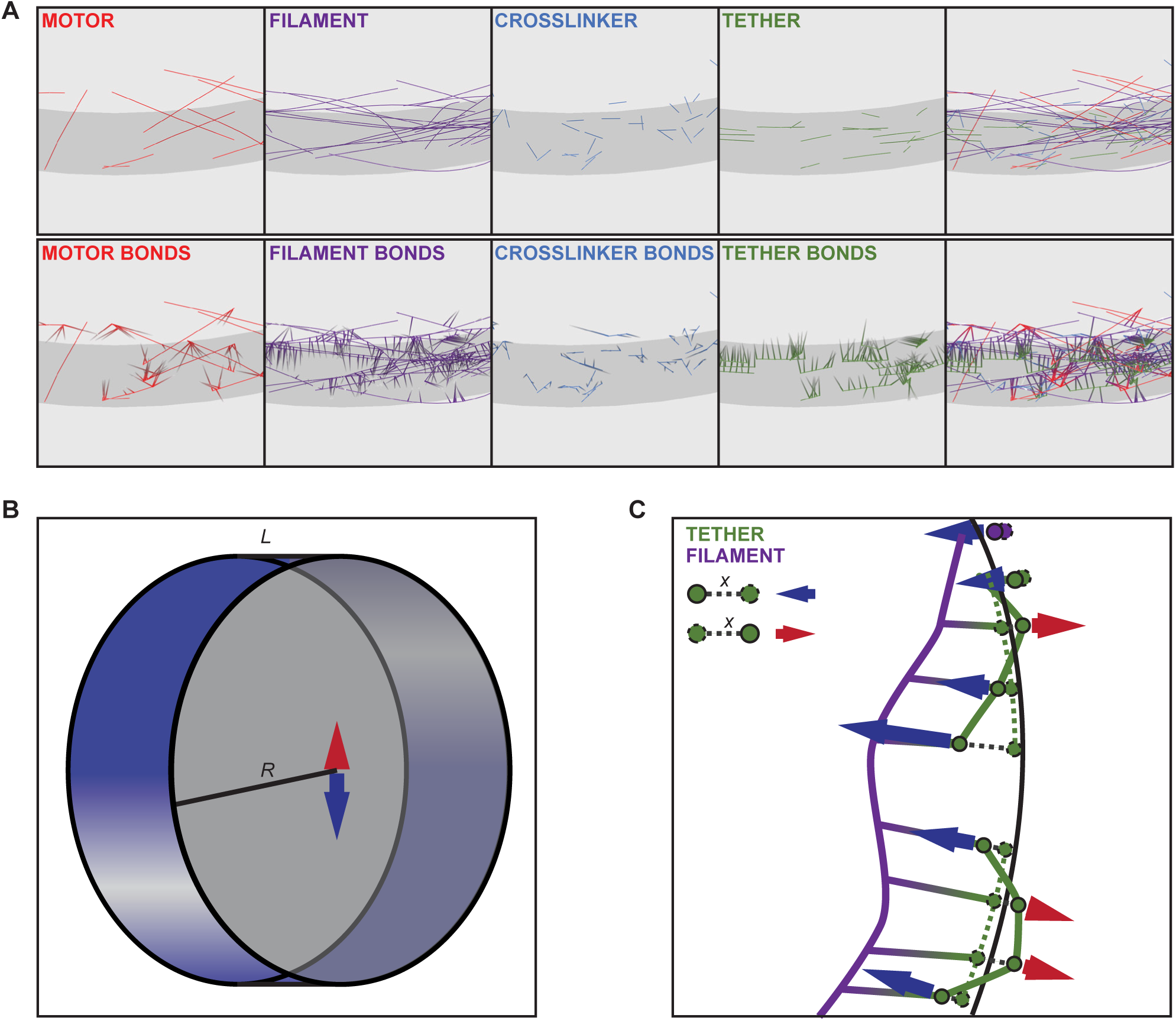
Three-dimensional contractile ring simulations. A) Individual components (top) and their bonds (bottom) shown through the section of simulated contractile ring shown in Figure 2. Tethers and their bonds appear the most closely associated with the dynamic space (gray surface). B) Representative schematic of the dynamic three-dimensional space with circumferential surface in blue and radial faces in gray. Arrows represent the sum radial forces of constriction (blue) and resistance to membrane deformation (red) that are calculated to generate constriction. C) Zoomed in schematic like D, but with a single filament linked to two tethering components (green) showing how displacement forces are calculated (blue arrows) along the points representing the tethering components. Solid dots represent actual positions of segment points, dashed dots represent their projection onto the space surface (black solid line). Solid lines represent actual positions of fibers, dashed lines represent their projections onto the space surface. Radial displacement, x, is calculated for all points to generate radial displacement force which can be contractile (arrows point left) or expansile (arrows point right) based on whether displacement is inside or outside the space surface. These forces are summed for all components to generate the resultant net radial force (single blue arrow in B).

**Figure S3.**
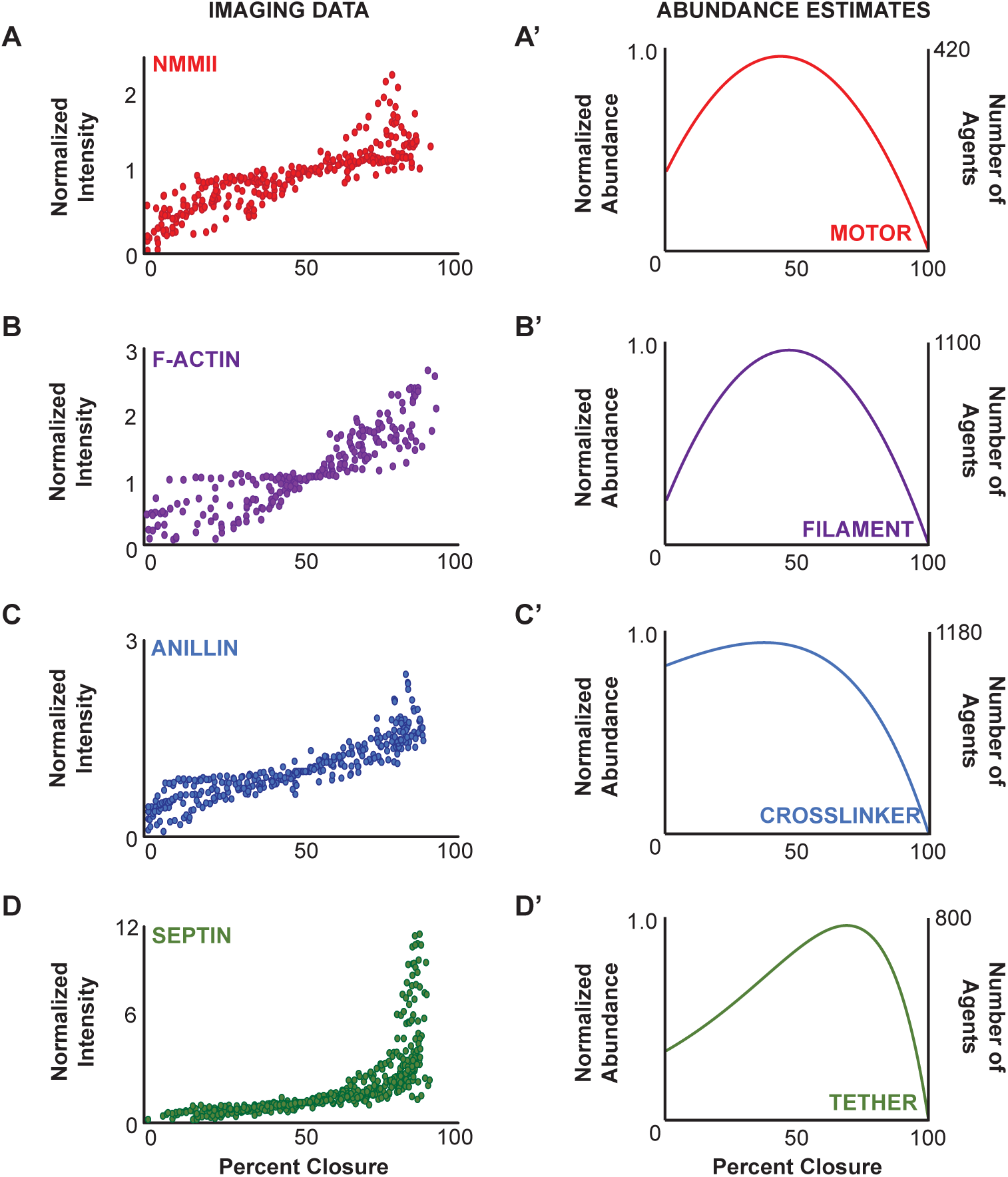
Calculation of abundance curves from biological datasets. A-D) Normalized intensity plots for average ring signal intensity of all cells of each imaged strain across all time points. A) NMMII fluorescent transgene (NMY-2::GFP). B) F-actin reporter fluorescent transgene (mKate::LifeAct). C) Anillin fluorescent transgene (mNeonGreen::ANI-1). A) Septin fluorescent transgene (UNC-59::GFP). Intensity for all was normalized so that the standard 1.0 would occur at 50% closure thus normalizing time and intensity to an internal metric. A’-D’) Normalized abundance curves generated for each of the contractile ring components based on measurements taken in A-D. Normalized abundance was estimated from best-fit lines for A and B and from best-fit exponential curves for C and D with maximums set at 1.0 (left Y-axis). Estimated number of agents as calculated from fission yeast cytokinesis literature (right Y-axis).

**Figure S4.**
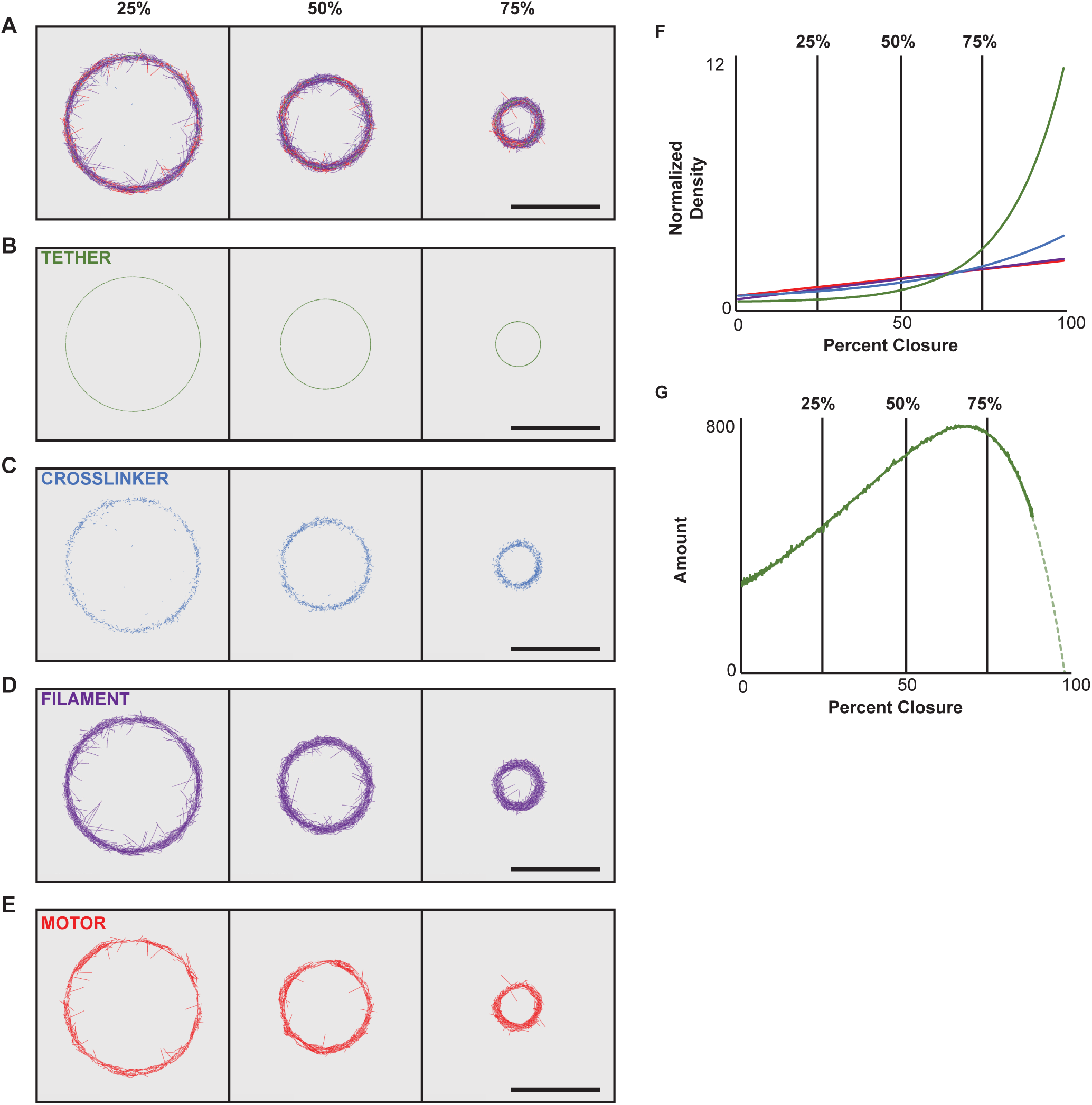
Component density throughout closure in simulated contractile rings. A-E) Simulation timepoints showing all components (A), tethers only (B), crosslinkers only (C), filaments only (D), and motors only (E) in two-dimensional whole ring projections at 25% closure (left column), 50% closure (middle column) and 75% closure (right column). F) Graph of normalized component density at all timepoints shown in A-E. Black lines along the graph represent the three timepoints selected for A-E. G) Graph of total tether component amount in a representative simulation (solid green) versus the calculated expected amount for a biological ring of the same size (dashed green). Scale bars = 15 μm.

**Figure S5.**
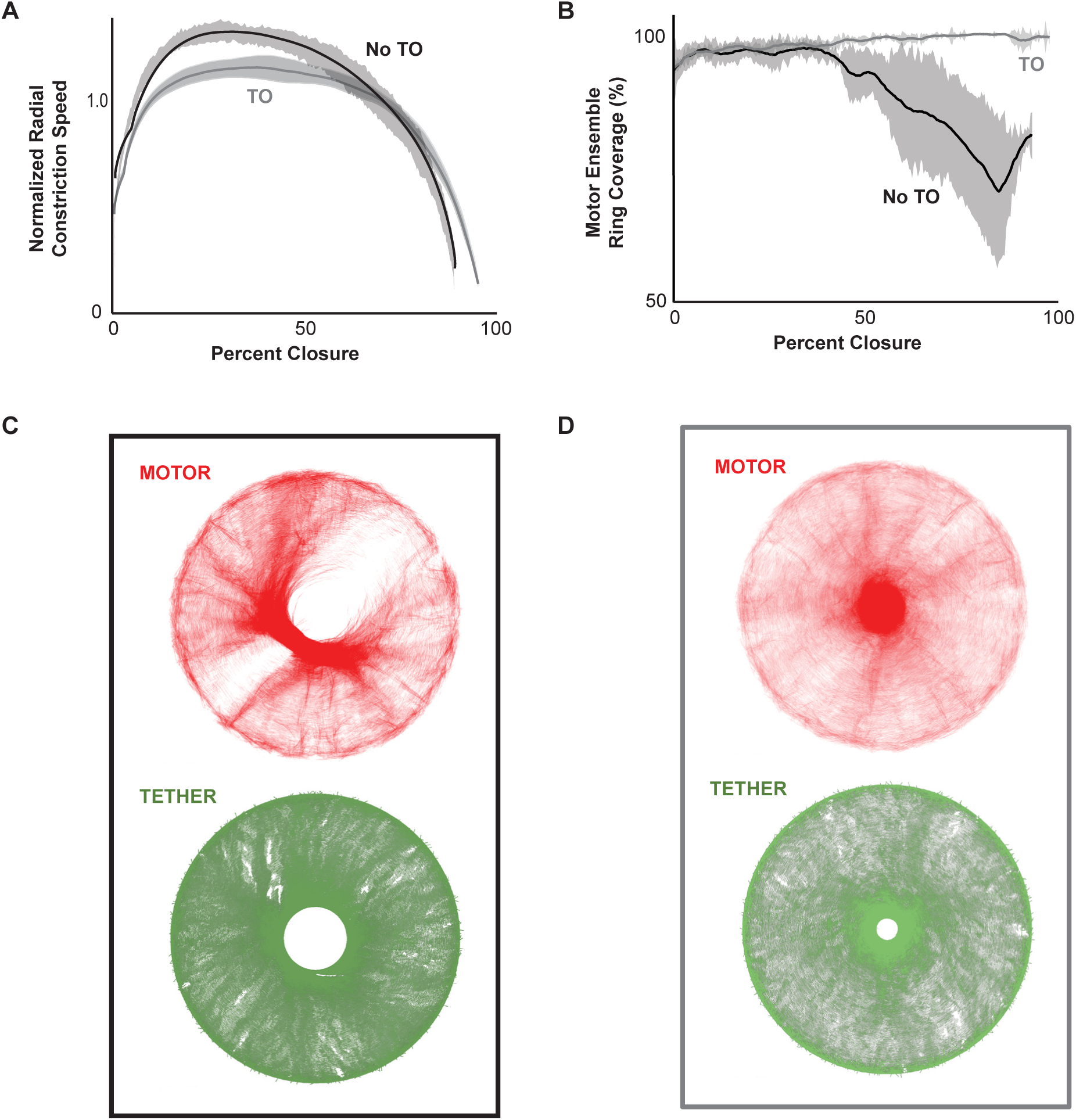
Turnover of components is required for contractile ring stability and full coverage. A) Graph showing closure dynamics for simulations with no component turnover (No TO; black) and with turnover of all components (TO; gray). Though both sets of simulations went beyond 90% closure on average 1/20 failed without turnover (dotted line) compared to 0/20 with turnover. B) Graph of motor component coverage of the simulated contractile rings with and without turnover. Without turnover motor component coverage of the contractile ring falls below 80% on average, with the simulation that failed to constrict fully falling below 60% (dotted line). With turnover motor distribution stayed above 90% ring coverage. C) Time-projection of motor (red) and tether (green) components throughout the simulation that failed constriction. Top panel shows motor clustering and an eventual full loss of coverage along a substantial portion of the ring (white space). Bottom panel shows consistent coverage of tethering components but a stall in constriction that results in incomplete closure. D)Time-projection of motor (red) and tether (green) components throughout a representative simulation with turnover. Both components show total ring coverage throughout the simulation (white space here is the result of contrast adjustment).

**Table S1.**
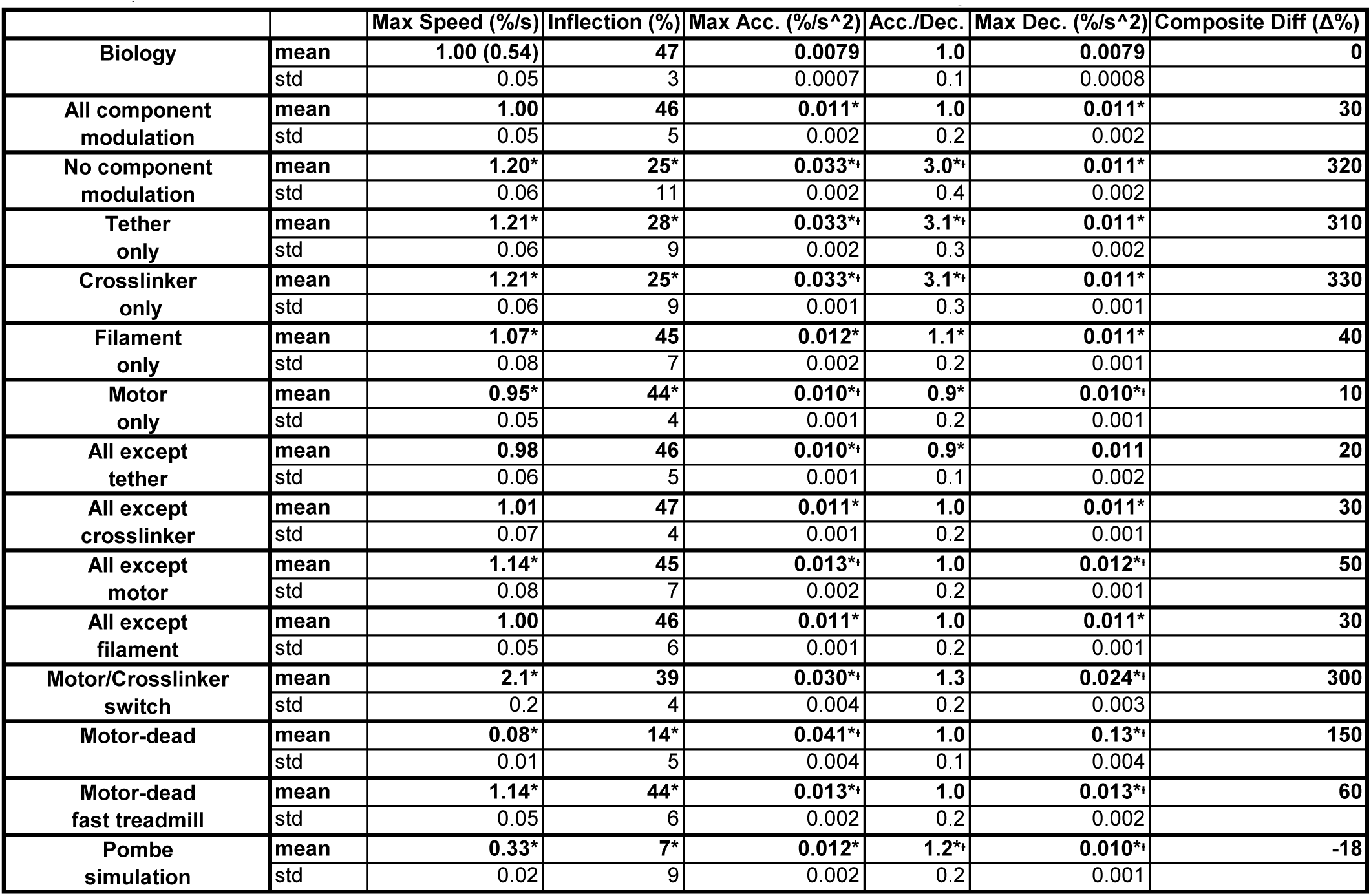
Quantification of constriction kinetics for in vivo and in silico contractile rings. Table shows the five metrics used to describe the constriction kinetics of our contractile ring data for all data. Max Speed: the normalized maximum speed, where biological/simulation standard max speed was set to 1.0. Inflection: describes at what point (as a % of initial ring size) maximum speed was reached. Max Acc.: the peak acceleration during the initial increase in constriction speed. Acc./Dec: the ratio between peak acceleration and peak deceleration. Max Dec.: the peak deceleration during the decrease in constriction that occurs after inflection. Composite Diff: describes the overall divergence of each simulation from the biological standard (first row) as a percent difference from the biological standard. * Denotes statistically significant difference by two-tailed t-test between the measurement and its biological equivalent (first row). ᶧ Denotes statistically significant difference by two-tailed t-test between the measurement and the simulation standard where all four main contractile ring components have dynamic abundance (second row). std rows for each sample show standard deviation from the mean; n = 23 for biological data (first row), n = 30 for all simulations (all rows after the first).

|  |  | Max Speed (%/s) | Inflection (%) | Max Acc. (%/s^2) | Acc./Dec. | Max Dec. (%/s^2) | Composite Diff (Δ%) |
| --- | --- | --- | --- | --- | --- | --- | --- |
| Biology | mean | 1.00 (0.54) | 47 | 0.0079 | 1.0 | 0.0079 | 0 |
|  | std | 0.05 | 3 | 0.0007 | 0.1 | 0.0008 |  |
| All component modulation | mean | 1.00 | 46 | 0.011* | 1.0 | 0.011* | 30 |
|  | std | 0.05 | 5 | 0.002 | 0.2 | 0.002 |  |
| No component modulation | mean | 1.20* | 25* | 0.033<sup>si</sup> | 3.0<sup>si</sup> | 0.011* | 320 |
|  | std | 0.06 | 11 | 0.002 | 0.4 | 0.002 |  |
| Tether only | mean | 1.21* | 28* | 0.033<sup>si</sup> | 3.1<sup>si</sup> | 0.011* | 310 |
|  | std | 0.06 | 9 | 0.002 | 0.3 | 0.002 |  |
| Crosslinker only | mean | 1.21* | 25* | 0.033<sup>si</sup> | 3.1<sup>si</sup> | 0.011* | 330 |
|  | std | 0.06 | 9 | 0.001 | 0.3 | 0.001 |  |
| Filament only | mean | 1.07* | 45 | 0.012* | 1.1* | 0.011* | 40 |
|  | std | 0.08 | 7 | 0.002 | 0.2 | 0.001 |  |
| Motor only | mean | 0.95* | 44* | 0.010<sup>si</sup> | 0.9* | 0.010<sup>si</sup> | 10 |
|  | std | 0.05 | 4 | 0.001 | 0.2 | 0.001 |  |
| All except tether | mean | 0.98 | 46 | 0.010<sup>si</sup> | 0.9* | 0.011 | 20 |
|  | std | 0.06 | 5 | 0.001 | 0.1 | 0.002 |  |
| All except crosslinker | mean | 1.01 | 47 | 0.011* | 1.0 | 0.011* | 30 |
|  | std | 0.07 | 4 | 0.001 | 0.2 | 0.001 |  |
| All except motor | mean | 1.14* | 45 | 0.013<sup>si</sup> | 1.0 | 0.012<sup>si</sup> | 50 |
|  | std | 0.08 | 7 | 0.002 | 0.2 | 0.001 |  |
| All except filament | mean | 1.00 | 46 | 0.011* | 1.0 | 0.011* | 30 |
|  | std | 0.05 | 6 | 0.001 | 0.2 | 0.001 |  |
| Motor/Crosslinker switch | mean | 2.1* | 39 | 0.030<sup>si</sup> | 1.3 | 0.024<sup>si</sup> | 300 |
|  | std | 0.2 | 4 | 0.004 | 0.2 | 0.003 |  |
| Motor-dead | mean | 0.08* | 14* | 0.041<sup>si</sup> | 1.0 | 0.13<sup>si</sup> | 150 |
|  | std | 0.01 | 5 | 0.004 | 0.1 | 0.004 |  |
| Motor-dead fast treadmill | mean | 1.14* | 44* | 0.013<sup>si</sup> | 1.0 | 0.013<sup>si</sup> | 60 |
|  | std | 0.05 | 6 | 0.002 | 0.2 | 0.002 |  |
| Pombe simulation | mean | 0.33* | 7* | 0.012* | 1.2<sup>si</sup> | 0.010<sup>si</sup> | -18 |
|  | std | 0.02 | 9 | 0.002 | 0.2 | 0.001 |  |

**Table S2.** Biophysical parameter list for contractile ring simulations. Table shows biophysical parameters, or estimations for all components included in contractile ring simulations.

| Parameter: Value | Comments | Reference |
| --- | --- | --- |
| **Simulation** |  |  |
| Space:<br>circle, 1.5 $\mu\text{m}$ radius | *S. pombe*-sized *C. elegans* embryo | |
| Viscosity: 1 N.s/m<sup>2</sup> | *C. elegans* embryo cytoplasm | (Daniels et al., 2006) |
| Temperature:<br>0.0042 pN.$\mu\text{m}$ | Room temperature | |
| Time-step: 1 ms | 1 ms for calculations of forces; 10 ms for visualization |  |
| Diffusion: set by Langevin | Diffusion is handled by the Langevin operator based on the length of the filament. |  |
| **Actin-like Filaments** |  |  |
| Length: $0.5 \pm 0.2 \mu\text{m}$ | Based on *S. pombe* length estimates | (Wu and Pollard, 2005; Courtemanche et al., 2017) |
| Rigidity: 0.06 pN.$\mu\text{m}^2$ | | |
| Barbed end growth speed: 0.01 $\mu\text{m/s}$ | | (Pollard, 1986) |
| Pointed end shrinking speed: 0.01 $\mu\text{m/s}$ | | (Pollard, 1986) |
| **NMMII-like Motors** |  |  |
| Length: 0.3 $\mu\text{m}$ | From *in vitro* reconstituted mammalian NMMII | (Billington et al., 2013; Nagy et al., 2013; Melli et al., 2018) |
| Rigidity: 0.2 pN.$\mu\text{m}^2$ | Estimated from flexibility of motors on filament backbone | (Billington et al., 2013) |
| Binding rate: 0.2 /s |  | (Stam et al., 2015) |
| Binding range: 0.08 $\mu\text{m}$ | within an order of magnitude of apparent motor head reach | (Billington et al., 2013) |
| Unbinding rate: 1.71/s | $k_0$ | (Stam et al., 2015) |
| Stall force: 3.85 pN | $F_d$ | |
| Unbinding relationship | $k_{\text{off}}(F) = k_{\text{off}}^0 [\alpha_{\text{catch}} \exp(-F x_{\text{catch}} / k_B T) + \beta_{\text{slip}} \exp(F x_{\text{slip}} / k_B T)]$ (Guo) | (Guo and Guilford, 2006; Stam et al., 2015; Cortes et al., 2020) |
| Max speed: 150 nm/s | $V_0$ | (Melli et al., 2018) |
| Force-speed relationship | $V = V_0 * (1 - F/F_{\text{stall}})$ | |
| Stiffness: 50 pN/$\mu\text{m}$ | Empirical | |
| Motors per ensemble: 30 | Simulated as a filament binding couple tether to the motor filament backbone | (Melli et al., 2018) |
| Diffusion: set by Langevin | Diffusion is handled by the Langevin operator based on the length of the filament backbone. |  |
| **Simple Crosslinker** |  |  |
| Length: 0.022 $\mu\text{m}$ | Empirical | |
| Rigidity: 0.1 pN.$\mu\text{m}^2$ | Empirical | |
| Binding rate: 10/s | Empirical |  |
| Binding range: 0.05 $\mu\text{m}$ | Empirical | |
| Unbinding rate: 0.3/s | Empirical |  |
| Unbinding force: 5 pN | Empirical |  |
| Stiffness: 50 pN/$\mu\text{m}$ | Empirical | |
| Diffusion: set by Langevin | Diffusion is handled by the Langevin operator based on the length of the filament backbone. |  |
| **Anillin-like Crosslinker** |  |  |
| Length: 0.04 $\mu\text{m}$ | Empirical | |
| Rigidity: 0.1 $\text{pN}\cdot\mu\text{m}^2$ | Empirical | |
| Binding rate: 10/s | Empirical |  |
| Binding range: 0.05 $\mu\text{m}$ | Empirical | |
| Unbinding rate: 0.3/s | Empirical |  |
| Unbinding force: 5 pN | Empirical |  |
| Stiffness: 50 $\text{pN}/\mu\text{m}$ | Empirical | |
| Diffusion: set by Langevin | Diffusion is handled by the Langevin operator based on the length of the filament backbone. |  |
| **Septin-like Tether** |  |  |
| Length: 0.1 $\mu\text{m}$ | | (Bridges et al., 2014) |
| Rigidity: 0.049 $\text{pN}\cdot\mu\text{m}^2$ | | (Bridges et al., 2014) |
| Binding rate: 10/s | Empirical |  |
| Binding range: 0.05 $\mu\text{m}$ | Empirical | |
| Unbinding rate: 0.1/s | Empirical |  |
| Unbinding force: 10 pN | Empirical |  |
| Stiffness: 50 $\text{pN}/\mu\text{m}$ | Empirical | |
| Diffusion: fast | Diffusion is not simulated. Instead, we assume that concentration of unbound crosslinkers is uniform |  |

